# Cascaded-CNN: Deep Learning to Predict Protein Backbone Structure from High-Resolution Cryo-EM Density Maps

**DOI:** 10.1101/572990

**Authors:** Spencer A. Moritz, Jonas Pfab, Tianqi Wu, Jie Hou, Jianlin Cheng, Renzhi Cao, Liguo Wang, Dong Si

## Abstract

Cryo-electron microscopy (cryo-EM) has become a leading technology for determining protein structures. Recent advances in this field have allowed for atomic resolution. However, predicting the backbone trace of a protein has remained a challenge on all but the most pristine density maps (< 2.5Å resolution). Here we introduce a deep learning model that uses a set of cascaded convolutional neural networks (CNNs) to predict Cα atoms along a protein’s backbone structure. The cascaded-CNN (C-CNN) is a novel deep learning architecture comprised of multiple CNNs, each predicting a specific aspect of a protein’s structure. This model predicts secondary structure elements (SSEs), backbone structure, and Cα atoms, combining the results of each to produce a complete prediction map. The cascaded-CNN is a semantic segmentation image classifier and was trained using thousands of simulated density maps. This method is largely automatic and only requires a recommended threshold value for each evaluated protein. A specialized tabu-search path walking algorithm was used to produce an initial backbone trace with Cα placements. A helix-refinement algorithm made further improvements to the α-helix SSEs of the backbone trace. Finally, a novel quality assessment-based combinatorial algorithm was used to effectively map Cα traces to obtain full-atom protein structures. This method was tested on 50 experimental maps between 2.6Å and 4.4Å resolution. It outperformed several state-of-the-art prediction methods including RosettaES, MAINMAST, and a Phenix based method by producing the most complete prediction models, as measured by percentage of found Cα atoms. This method accurately predicted 88.5% (mean) of the Cα atoms within 3Å of a protein’s backbone structure surpassing the 66.8% mark achieved by the leading alternate method (Phenix based fully automatic method) on the same set of density maps. The C-CNN also achieved an average RMSD of 1.23Å for all 50 experimental density maps which is similar to the Phenix based fully automatic method. The source code and demo of this research has been published at https://github.com/DrDongSi/Ca-Backbone-Prediction.

## I. Introduction

Proteins perform a vast array of functions within organisms. From molecule transportation, to mechanical cellular support, to immune protection, proteins are the central building blocks of life in the universe [1]. Despite each protein being composed from a combination of the same 20 naturally occurring amino acids, a protein’s functionality is mainly derived from its unique three-dimensional (3D) shape. Therefore, learning the details of a protein’s 3D structure is a prerequisite to understanding its biological function.

### A. Cryogenic Electronic Microscopy (Cryo-EM)

Currently, one of the leading techniques for determining the atomic structure of proteins is cryo-electron microscopy (cryo-EM). Cryo-EM is a relatively new technique which uses a high-energy electron beam to image vitrified biological specimens. In the past five years, more than 1,000 protein structures have been imaged at 4Å resolution or better in the EM databank using cryo-Electron Microscopy (cryo-EM). Among them, many are of detergent-solubilized membrane proteins [2] [3] [4] [5] [6] [7]. These high-resolution images make it possible to produce atomic level 3D models from the density maps.

### B. Protein Backbone Structure

From clean, high-resolution EM density maps (< 5Å) it is possible to distinguish the backbone structure of a protein [8] [9] [10]. A protein’s backbone is a continuous chain of atoms that runs throughout the length of a protein, see Fig. 1A. The backbone structure consists of a repeated sequence of three atom (carbon, nitrogen, alpha-carbon). Of these three atoms, the alpha-carbon (Cα) is particularly important as it is the central point for each amino acid residue within the protein. Therefore, predicting not only a protein’s backbone but also the locations of each Cα along that backbone can help determine where specific amino acids are located throughout the protein structure.

**Fig. 1.**
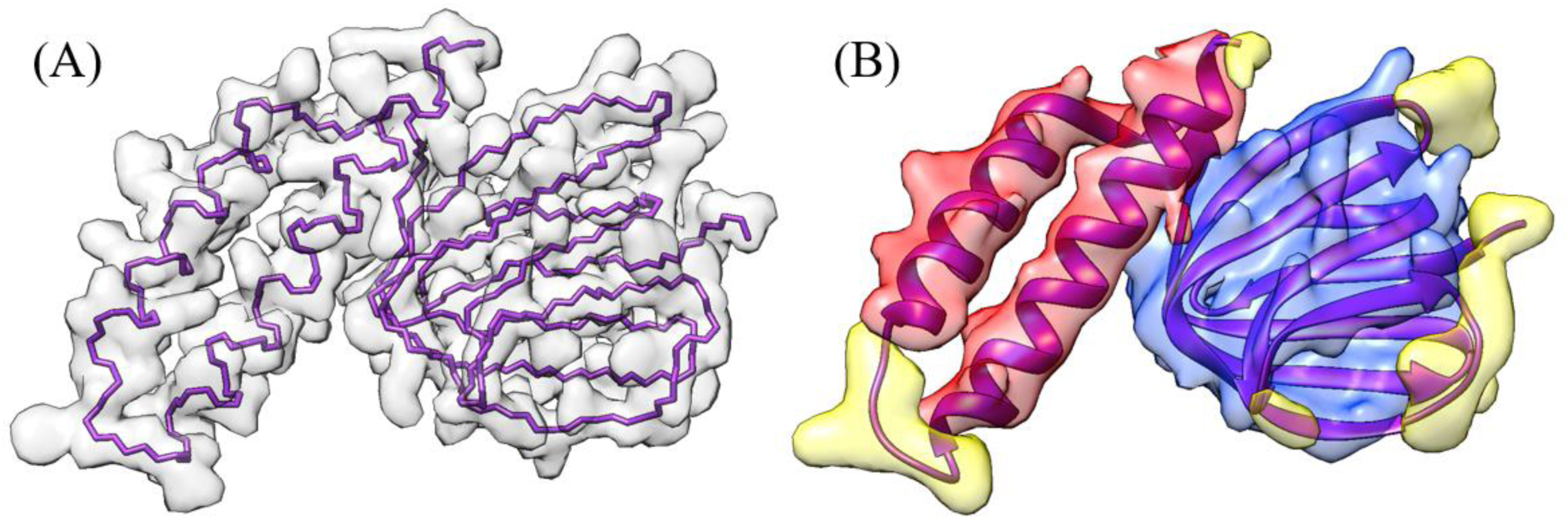
Simulated Density Maps from protein 1aqh at different resolutions. (A) shows a high-resolution map with underlying backbone trace. (B) shows a medium-resolution map with underlying ribbon structure. α-helix structures are colored red, β-sheets are colored blue, and the loops/turns are colored yellow.

### C. Protein Secondary Structure Detection

In addition to the backbone features of a protein, some of the most visually dominate features of cryo-EM density maps are the secondary structure elements (SSEs), see Fig. 1B. The three SSEs are α-helices, β-sheets, and turns/loops. At medium resolution, α-helices appear as long cylinders with a radius of approximately 2.3Å. β-sheets consist of multiple parallel beta strands that connect laterally by hydrogen bonds. While only distinguishable at 6Å resolution or better, β-sheets appear as flat or slightly wavy sheets. Turns/Loops are the final SSE. They occur in locations where the polypeptide chain of the protein reverses its overall direction. When imaged with cryo-EM, turns/loops often appear faint due to their relatively low electron density. This makes them one of the most challenging SSE to classify.

There are many methods for identifying SSEs at medium resolutions [11] [12] [13] [14] [15] [16] [17]. However, at higher resolution (< 2.5Å), the classic α-helix and β-sheet structures are not easily recognizable to the human eye. This is due to the large number of side chains that protrude off the backbone chain in high-resolution data. This makes predicting the SSEs at high-resolution potentially more difficult than at medium resolution.

### D. Current Protein Prediction Models

Ever since the first experimental density maps were released for protein structures, researchers have been developing software models to predict the various structural elements from each map. Some of the leading software models are now able to predict the atomic structure of a protein from its electron density map.

Phenix is a widely used molecular prediction software suite that has often been used in research since its initial release in 2010 [18]. A recent 2018 paper introduced a new molecular prediction method that combined the Phenix prediction software along with advanced post-processing techniques [19]. This method, henceforth referred to as the Phenix method, produced some of the most-complete prediction models. As a result, we used this method as a metrics benchmark for this research.

The Phenix method is a fully-autonomous prediction method which only requires a density map and a nominal resolution value as input. This method first sharpens the input density map using an automated map sharpening algorithm which aims to maximize the connectivity of high-density regions [20]. Then, for each part of the structure, various atomic models are generated using several independent prediction models, including one for SSEs and one for backbone tracing, among others [21] [22] [23]. The results from these predictions are ensembled and used to produce an initial predicted structure. This structure is then refined using any symmetry that is present in the protein. The Phenix method was tested on 476 experimental density maps and has, to date, produced the most complete prediction maps. This method also uses a unique set of metrics to measure the effectiveness of the prediction method. The RMSD method uses a one-to-one mapping of predicted to ground-truth Cα atom but only includes atoms that are within 3Å of the ground truth model. To measure prediction model completeness, the Phenix method calculates the percentage of matching Cα atoms between the predicted and ground-truth model within the same 3Å space. We use the same metrics when evaluating our deep-learning prediction technique.

RosettaES is a protein modeling software tool first developed at the University of Washington. RosettaES employs a modeling technique which consists of two general components: conformation sampling and energy evaluation [24]. Conformational sampling uses well-established physical characteristics of molecular structure as guides for model prediction. Examples of such characteristics include: the common torsion angles of atoms in the backbone structure, or the radius of α-helix secondary structures. Each of these structures has a very narrow band of potential values making them excellent constants to use when modeling protein structure. The energy evaluation process calculates the total energy of a predicted protein based on each predicted atom position along with each bonding angle between them. This value is compared to the expected lowest-energy state, which can be calculated from sequence information. Given that the lowest-energy state is likely closest to the native state of the protein, slight adjustments are made to the predicted protein structure to minimize the energy within its atomic structure thereby optimizing the prediction map for the protein.

Another leading backbone prediction model is the MAINMAST algorithm developed by researchers at Purdue University [25]. MAINMAST produces a backbone trace, consisting of a set of Cα atoms, from high density regions of an electron density map. This algorithm first identifies regions of high-density (high-density points are likely to be backbone structure) using mean shifting and then transforms them into a minimum spanning tree (MST). A Tabu search algorithm is applied to find a few thousand possible MSTs. For each MST, the amino acid sequence is mapped on the longest path in the tree using areas of high density as likely Cα atom locations. Each MST is rated based on the best fit. The highest scoring tree is chosen as the final prediction of the model.

In designing our experimental method, we leveraged techniques from each of these leading prediction methods. We employed a new conformational sampling technique similar to the RosettaES method. Our technique used constants such as: standardized distance between Cα atoms, mean α-helix radius, and common torsion angles between backbone atoms. Using these constants, we also invented a new Tabu-search scoring algorithm, similar to the one used in the MAINMAST method. Our Tabu-search was primarily used as a backbone path-walking algorithm. Finally, we employed the multi-prediction model approach of the Phenix method by creating a different CNN to predict the SSEs, backbone, and Cα atoms of each density map before stitching them together to form a final prediction map.

### E. Deep Learning Semantic Segmentation

This research aimed to use deep learning to create a predictive model capable of detecting the SSEs, backbone structure, and Cα atoms from electron density maps. The field of deep learning has proven to be very successful in the fields of image recognition and image classification [26] [27] [28]. This research used a specific image classification method known as semantic segmentation. With semantic segmentation each 2D-pixel or 3D-voxel of an image is classified independently rather than the entire image as a whole.

Until recently, semantic segmentation was accomplished through patch classification. Patch classification takes a slice of the input data and runs it through a convolution neural network (CNN). However, patch classification only classifies the center pixel of each patch meaning that the CNN would have to process a new patch for each pixel in the image. This technique is preferred when computing resources are limited because processing a small patch is much less computationally expensive than processing a full image.

However, with the recent advances in GPU technology, fully connected end-to-end networks are now able to perform semantic segmentation on full images in one pass. In 2014, research at UC Berkley used a Fully Convolutional Network (FCN) to perform semantic segmentation on the PASCAL-Context dataset [29]. Their method used an encoder-decoder architecture that removed the need for patch classification by essentially combining the calculations of the overlapping patch regions into a single end-to-end network. In 2015, the network *Segnet* aimed to improve the encoder-decoder architecture by forwarding the max-pooling indices from the encoder layer to the decoder layer to prevent the loss of global information in the image [30]. Later that year, researchers at Princeton University used a technique called dilated convolution which made it possible to perform semantic segmentation without the encoder-decoder architecture [31]. Dilated convolutions are preferred when a convolution needs to increase the field of view without reducing the resolution of the image.

In this paper, we leveraged the architecture of each of these semantic segmentation classifiers. Previous deep learning methods that were used for SSE prediction used patch classification [12]. Our model levered the Fully Connected Network design to eliminate the need for patch classification and instead use semantic segmentation to classify a full 3D image in a single pass. It also used data forwarding, inspired by Segnet, to allow for segregated learning. Finally, this model used dilated convolutions to increase the field of view while maintaining the input image dimensionality.

## II. Methods

### A. Data Collection/Generation

Predictive models are only successful if they are trained with representative data. Since, high-resolution cryo-EM maps are still somewhat scarce we trained our model using simulated cryo-EM maps instead of experimental maps. This design decision allowed us to save all the experimental maps for verification.

Simulated cryo-EM maps can be generated from protein databank (PDB) files^1^ using a script from the EMAN2 package called *pdb2mrc* [32]. This script takes each atom in a PDB file and produces a 3D Gaussian electron density. It then sums the Gaussian density of all the simulated atoms on a 3D grid to produce a complete electron density map for the entire protein. This simulation method produces electron density maps that are very representative of their experimental counterparts with the primary difference being that a simulated map has no experimental inaccuracies.

To produce a large enough dataset for training, we used over 7,000 PDB files to generate simulated density maps using the *pdb2mrc* script. Each map was simulated at a different resolution to produce a higher amount of variance in the training set and prevent overfitting in the model. In addition to simulating the full PDB structure, we also had to simulate the labeled data for each map. This step involved editing the PDB file to retain only atoms that were part of the predicted structure and running *pdb2mrc* to produce a labeled map for each structure being predicted. When selecting PDB files, we selected maps with a sequence structure that was at least 50% unique relative to all other maps in the training set. This ensured that our training data was diverse and well-representative of the large range of protein structure found in nature.

The data generation pipeline is outlined in Fig. 2. The output of this pipeline was two HDF5 files: one training file and one testing file. The training file consisted of approximately 24,000 simulated density maps^2^ along with the corresponding SSE, backbone, and Cα labeled maps for each protein. The testing HDF5 file, which was generated to measure the accuracy of the neural network as it trained, consisted of 1,024 maps along with the same configuration of labeled maps. All testing maps were unique, and no map rotation was performed to increase the testing data size.

**Fig. 2.**
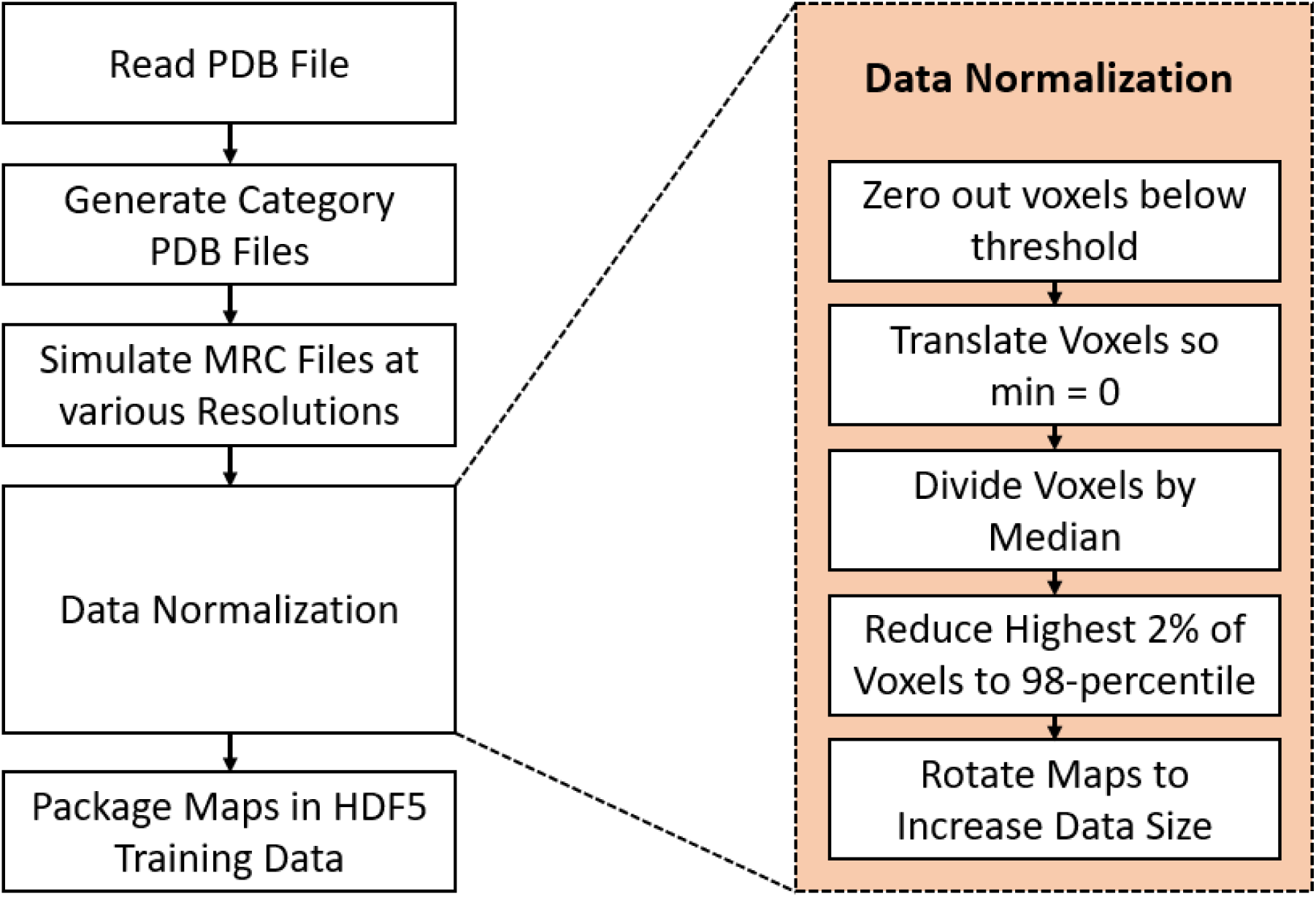
Simulated Data Generation Pipeline including details about the data normalization process. The output of this pipeline was an HDF5 file containing all the data used to train the prediction model.

To ensure uniformity among each electron density map, extensive data normalization was used to produce a common input data format. There were five data normalization steps, see Fig. 2. First, all voxels with an electron density less than a resolution dependent threshold were zeroed out^3^. This removed low intensity areas of the simulated maps which differed significantly from their experimental counterparts. It also allowed the neural network to exclusively train with voxels of high-intensity, which are often more representative of protein structures. After this step all voxel intensity values were reduced by a threshold value so that the spectrum of electron densities within each map started at zero. The third step of data normalization involved dividing all voxels by the median voxel value in the electron density map. This step normalized the voxel values and ensured that each map had a similar data range. After this, data outliers were removed by capping all voxels at the 98th-percentile voxel intensity. Finally, each training map was copied and rotated by a varied angle to increase the total training data size.

### B. Cascaded Convolutional Neural Network

Building off the previous semantic segmentation convolutional neural networks, we designed a cascaded convolutional neural network (C-CNN) consisting of three feedforward dilated neural networks. The high-level architecture is shown in Fig. 3. This design allowed us to train all the neural networks simultaneously. The input to the C-CNN was a 64×64×64 tensor representing the 3D electron density of a protein. However, because density maps vary greatly in size across each dimension, an extra step was required before the model could process maps of a different size. Each map was split into 64×64×64 cubes that overlapped by 7 voxels on each face. Each cube was evaluated by the C-CNN independently and then the resulting output cubes were stitched back together to reconstruct the full image. However, only the center 50×50×50 voxels were used to reconstruct the image. At each face the 7-voxel overlap region was disregarded. This method allowed us to process density maps of any size without losing spatial information at each cube’s boundary.

**Fig. 3.**
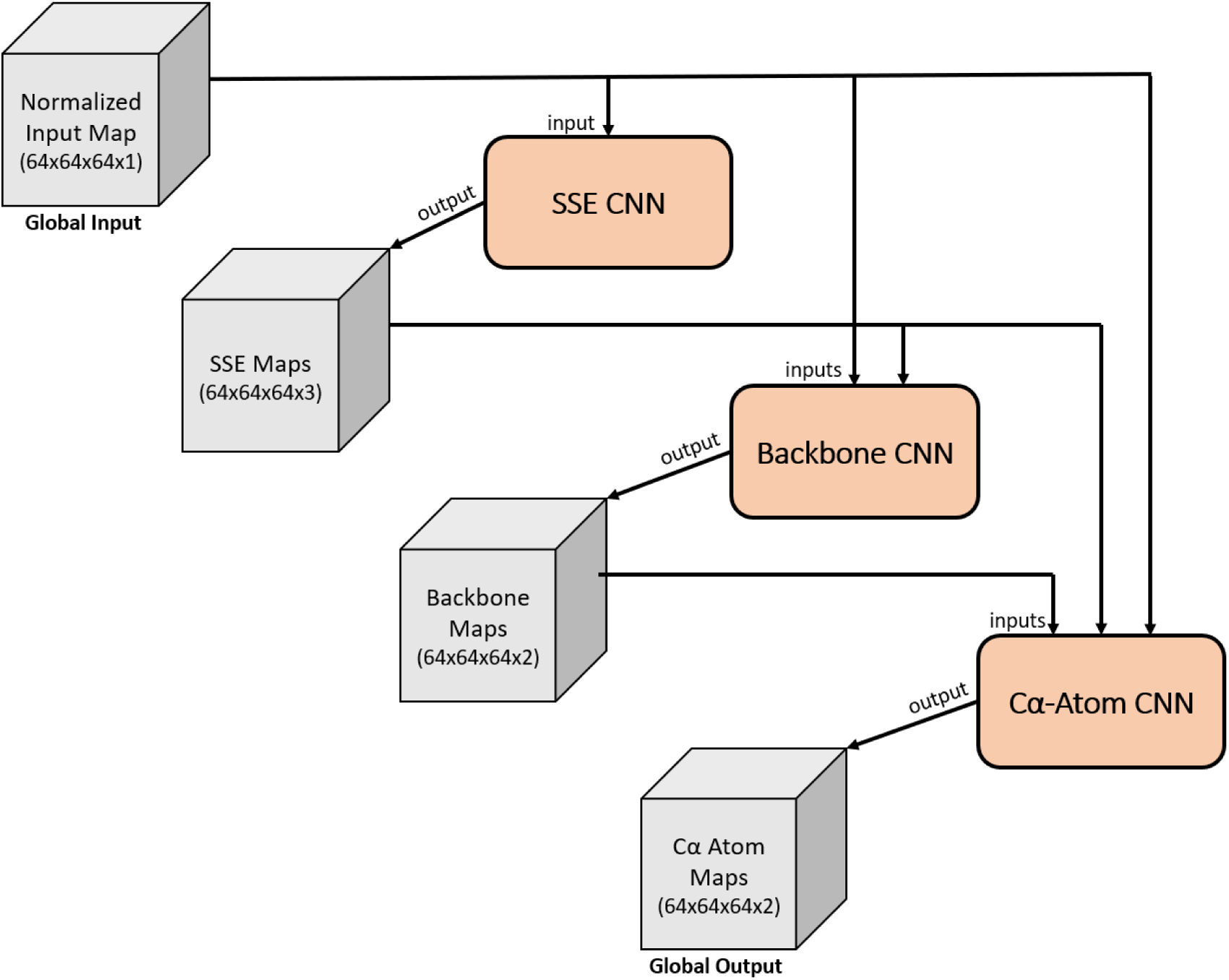
Cascaded Convolutional Neural Network. The input/output of each stage is shown as a gray cube with the given dimensions. Each CNN is represented by a tapered salmon rectangle. Results from each CNN are forwarded along with previous input data to the next CNN.

Inside the C-CNN, the input map was forwarded to each of the three neural networks. The first network was the SSE CNN. It predicted voxels as α-helices, β-sheets, or loops/turns and output a confidence map for each SSE. The three SSE maps along with the input electron density map were forwarded to the backbone CNN which produced two confidence maps representing whether each voxel was part of the backbone structure of the protein or not. The final CNN in the C-CNN was the Cα-Atom CNN. This network took all the previous maps and produced two output maps representing the confidence of a voxel being part of a Cα atom or not.

### C. Convolutional Neural Network Architecture

The three neural networks were very similar, each having the same number of layers and same type of layers. Their detailed structure is illustrated in Fig. 4. Each neural network had four layers. The first and fourth layers were regular 3-D convolutional layers with a stride of one. The second and third layers were dilated convolutions, also with a stride of one. Each dilated convolutional layer used a dilation rate of two. Following each dilated convolution was a leaky ReLU activation function. A leaky ReLU was preferred over a standard ReLU activation function to improve back-propagation and reduce the problem of vanishing gradient decent. Dilated convolutional layers were used to increase the receptive field while maintaining the image size. This is crucial for semantic segmentation because it maintains a one-to-one ratio of input voxel to output voxel.

**Fig. 4.**
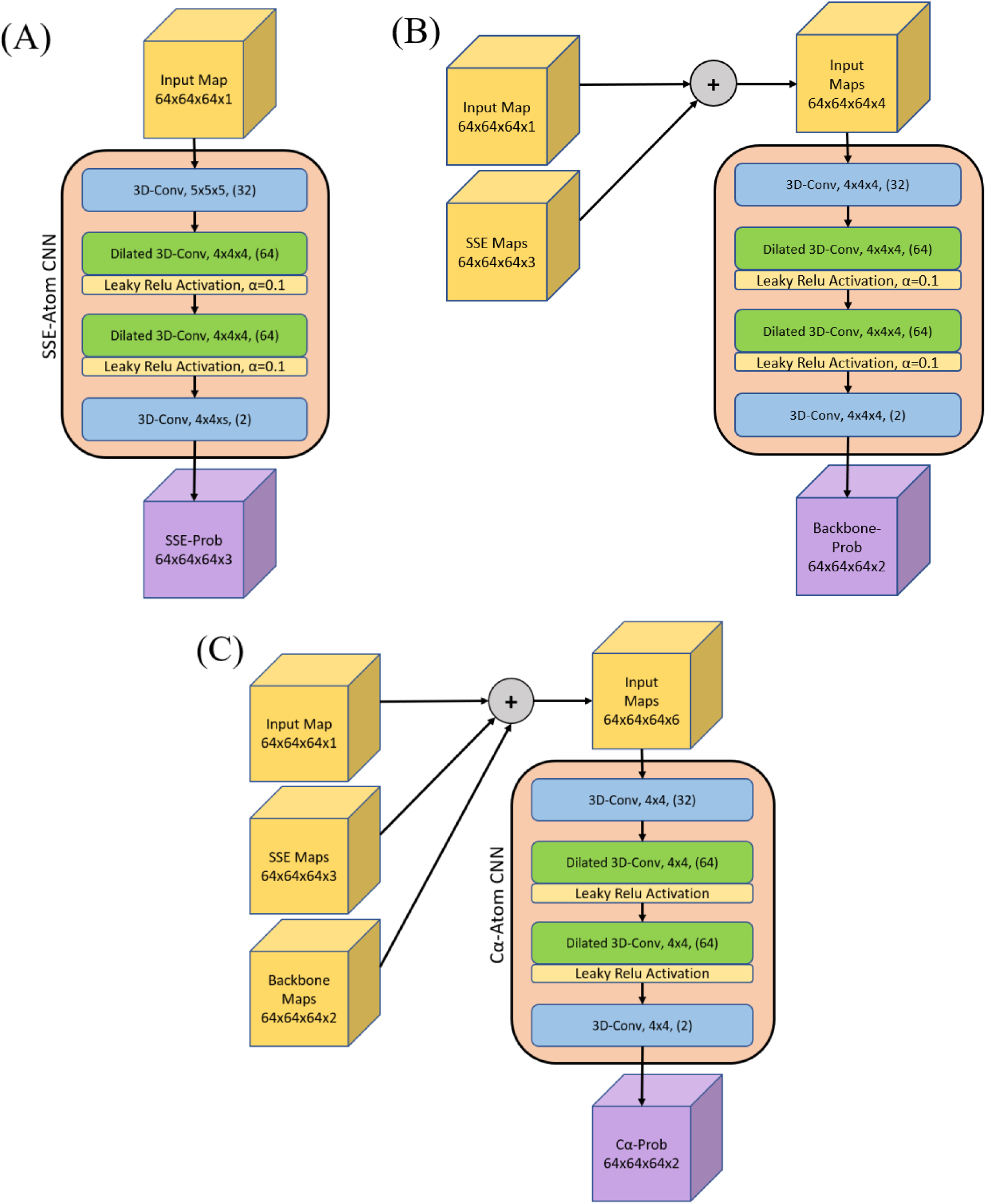
Detailed architecture of each of the 3D convolutional neural networks (CNN). (A) contains the Secondary Structure Elements (SSE) CNN. (B) contains the Backbone CNN. (C) contains the Cα-Atom CNN. Each CNN, including all its layers, are shown within the salmon colored boxes. The input to each CNN is noted by the yellow cubes. The Backbone CNN (B) and the Cα-Atom CNN (C) take input that is a contamination of various maps. The size of each input is noted by the dimensions listed at the base of each input cube. Each layer in each CNN is denoted by its name/function, kernel size (NxNxN), and finally the output number of filters (inside parenthesis) for that layer. Each leaky ReLU activation function used an alpha value of 0.1. The output of each CNN is noted by the purple cube. The dimensions for each output are listed at the base of the cube.

Each of the three neural networks used the same number of filters per layer for the first three layers: 1st layer: 32 filters, 2nd layer: 64 filters, and 3rd layer: 64 filters. The 4th and final layer differed for each network, but it was always equal to the number of output classes. Using more filters usually leads to higher accuracy. However, even a small increase from these numbers greatly slowed the network training. Therefore, we settled with these values as it was an optimal compromise between accuracy and speed. Other than the slight difference in filters per layer, the only other difference among the networks was a small difference in kernel size in the standard convolutional layers. The kernel size was larger (5×5×5 vs. 4×4×4) in the backbone and SSE CNNs to account for the need for a larger receptive field to better predict those structural elements.

### D. End-To-End Model Pipeline

The cascaded convolutional neural network is only a piece of the full backbone prediction model. The full model is shown in Fig. 5. The primary input to the full model was an MRC file or MAP file containing a 3D tensor of the electron density of the protein. The only other input was a manually selected threshold value which was used to zero out low intensity voxel values^4^. Selecting a proper threshold is challenging because each density map is very different in intensity. However, the recommended contour level on the EM Databank website^5^ is a good value to start with. The final output of the entire prediction model was a PDB file containing a set of traces where each trace is a set of connected Cα atoms. This PDB file also contains SSE labels for each Cα atom. These labels are determined from both the SSE output maps and the helix-refinement. Using the final output map the RMSD and the Percentage of Matched Cα atoms metrics were calculated.

**Fig. 5.**
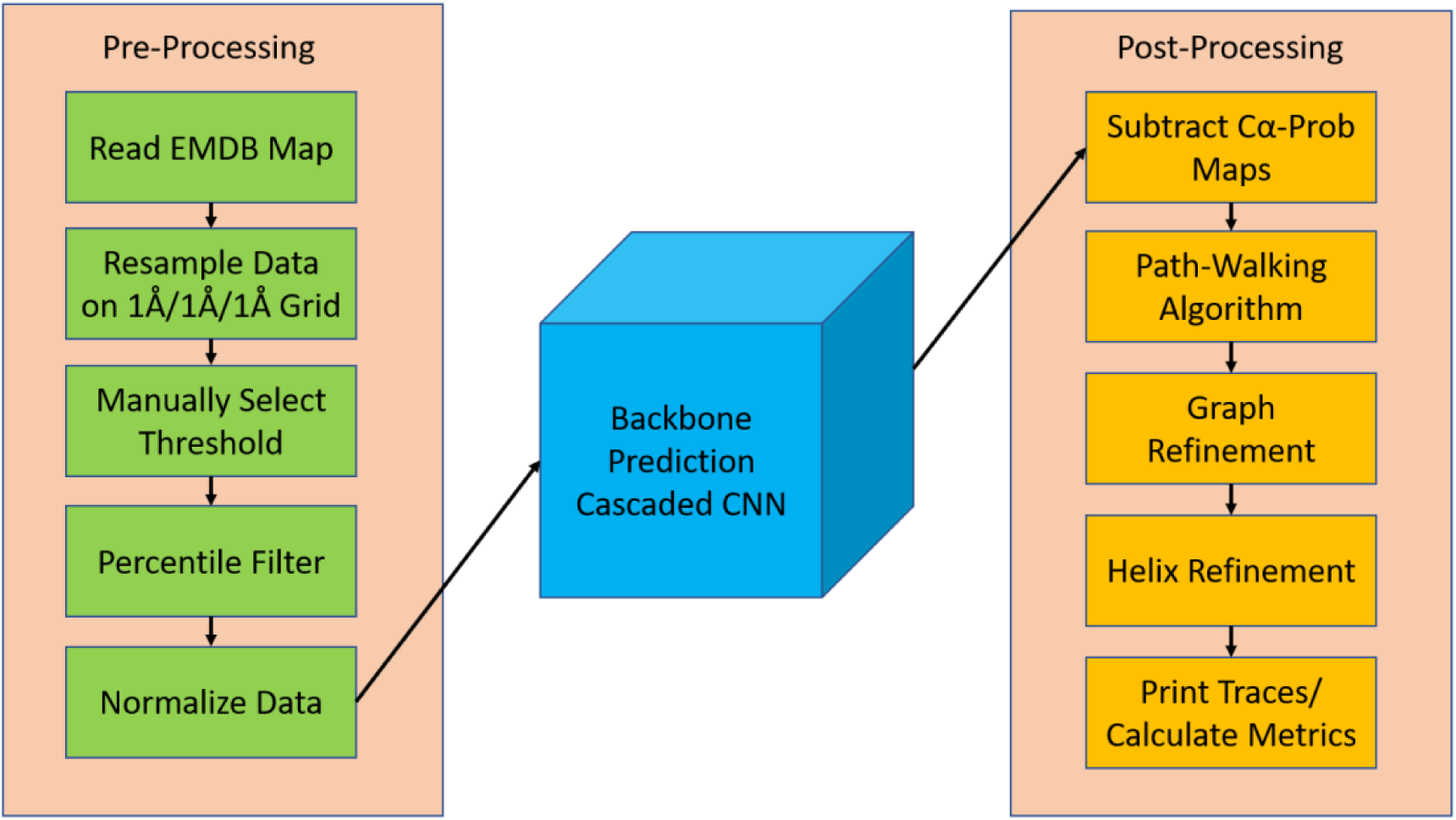
Full Backbone Prediction Model. Model includes data preprocessing, the cascaded convolutional neural network (after training), and post-processing.

### E. Pre-Processing

The goal of pre-processing experimental density maps before sending them into the cascaded convolutional neural network is to make them as similar as possible to the simulated maps that the C-CNN was trained with. Unlike simulated maps, experimental maps have large variance in local classified resolutions, electron density values, and molecular shapes. This can be attributed to the wide range in flexibility of biological molecules, cryo-EM imaging devices, different experimental procedures, and small natural artifacts that appear as part of the cryo-EM imaging process. Combined, these issues make experimental maps difficult to normalize.

The first step of preprocessing was to remove any noise or irrelevant electron density data. This step was accomplished by zeroing out all voxels that were greater than 6Å away from the ground truth structure of the protein map. Once cleaned, the density map was resampled so that each dimension (x, y, z) had a voxel-size of exactly 1Å. There is a wide range of voxel-sizes for each experimental map and many often have a different value for each axis. Therefore, resampling was crucial because the C-CNN was trained with simulated maps that had a voxel-size of exactly 1Åx1Åx1Å. This step was easily accomplished by using the UCSF Chimera tool along with the internal Chimera command vop resample.

After resampling, the new map was preprocessed using the same method as outlined in Fig. 2 with the only difference being that the threshold was manually selected for each density map.

### F. Network Output

After the C-CNN processed the input map it produced confidence maps for the protein’s SSEs, backbone, and Cα atom locations. The output maps for EMDB-8410 (chain-A) are shown in Fig. 6. Each voxel in the map was assigned a confidence value by the network. The final classification of a voxel was determined by the max voxel value of each of the output maps for a given neural network.

**Fig. 6.**
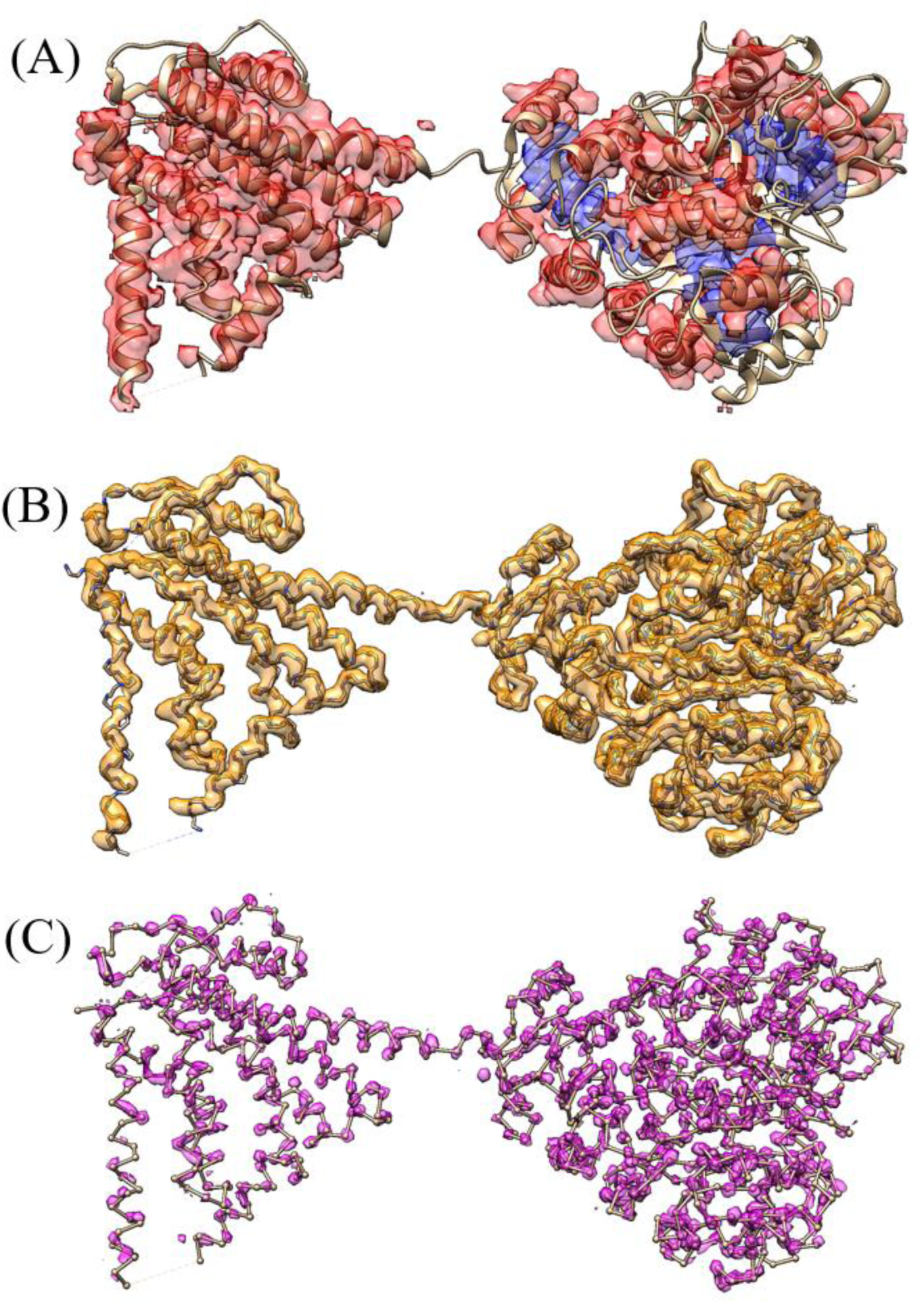
Confidence Map Output for EMDB-8410 (Chain-A). (A) is the combination of the α-helix and β-sheet prediction map after applying the max function, (loops/turns map omitted for readability). (B) is the backbone confidence map (>40% confidence) with the ground truth backbone structure shown for reference. (C) is the Cα-Atom confidence map (>50% confidence) with the ground truth ball and stick representation of EMDB-8410 shown for reference.

### G. Path-Walking Algorithm

Although the C-CNN assigns confidence values to specific features of the protein, post-processing algorithms were required to piece together that information and generate a final prediction trace. This was accomplished with a path-walking technique that processed the confidence maps to produce a final PDB file that contained exact Cα atom locations along the protein’s backbone.

The path-walking technique walked through high-confidence areas of the backbone map and connected areas of high Cα atom confidence using a novel tabu-search algorithm designed specifically for this research. The tabu-search algorithm scored each potential future movement based on a location’s local density prediction confidence and distance. Additionally, it also incorporated the backbone atom torsion angles and common radius of α-helix secondary structures as weights when finding the optimal next Cα atom.

The path-walking algorithm walked until it either reached an area of the protein that had already been processed or until it reached an area of the protein where no more suitable Cα atoms could be found. Upon reaching the end of a single trace, the path-walking algorithm would search the Cα confidence map for any other areas of the protein that might contain additional untraced Cα atoms and, if found, would walk each additional trace. This process was repeated until all high-confidence areas of the Cα prediction map had been explored. The output of the path-walking algorithm was a PDB file consisting of a series of disconnected traces where each trace contained a chain of Cα atoms.

### H. Graph Refinement

The disconnected traces from the path-walking algorithm represented partial backbone traces in the protein. However, there were many false positive traces that were the result of side chains and shortcuts between backbone structures being incorrectly classified as backbone traces. To remedy this issue, two refinement steps were required to improve the predicted traces: path combination and backbone refinement. In order to complete these two steps, the backbone traces were converted from a list of Cα locations and connections into a graph where each Cα atom was represented by a node and each connection to another Cα atom was represented by an edge.

The goal of the path combination step was to combine a set of disjoint graphs (formerly traces) into a fully connected graph that is more representative of the protein’s backbone structure. We used a depth first search to walk from any given Cα atom within a disjoint graph to both end points of that trace. Using these endpoints, this algorithm would then examine all other Cα nodes in the protein graph to determine if another Cα atom was within 3Å of the end point Cα atom. If it was, then the end point Cα atom’s location was reassigned to be equal to the other Cα atom. This process helped combine neighboring disjoint graphs (traces) into a fully connected graph.

After path combination, the fully connected graph resembled a protein’s backbone structure, but it still had many side chain and backbone trace shortcut connections. These false-positive connections meant that many Cα nodes in the graph had three or even four edge connections to other Cα atoms. The next step in the graph refinement process was to remove the false-positive connections so that the remaining graph only contained true-positive Cα node and Cα edge connections. This refinement process was broken down into three steps: side chain removal, loop removal, dead-end point removal.

Side chain removal involved examining every Cα node in the graph to determine if it might be part of a false-positive side chain connection. If a node had three or more edges (trinary node), it was likely that one of the three edges was a side-chain connection. This algorithm would use a depth first search (DFS) to walk along each of the three paths leading from a trinary node and stop once it found either an ending node (only one edge), or another node with three or more edges. It would compare the total depth reached for each of the three DFS edge walks. If one path had a depth of three or less while the other two paths both had depths greater than the shortest path, then the shortest path was considered a side chain connection and removed from the graph. This algorithm proved to be very effective at removing side chain and false-positive shortcut connections between true parallel backbone traces.

After removing the side chains from the fully connected graph, it was necessary to remove small loops within the graph. These loops were the result of false-positive shortcut traces within α-helix elements of the protein. The goal of this method was to remove the false-positive half of each loop leaving the true backbone structure in place. The approach was similar to side chain removal. It would find any Cα nodes that contained three or more edges and then path walk each trace until it reached an end-node or trinary connection. However, in this case, if two paths terminated in the same trinary node then the combined two paths were considered a loop. To remove the false-positive side of the loop, this method would calculate the density along each path using a 1Å radius cylinder and remove the path with the lower average density value. This approach made the assumption that the backbone structure of the protein had a higher density than another false-positive path.

The final step of the graph refinement process was to remove dead-end nodes. These resulted from side-chains that did not connect to another backbone trace of the protein but did nonetheless protrude off the true backbone structure. Removing these was accomplished by finding all trinary Cα nodes in the graph and then walking down each trace extending from that node. If any path had a depth of two or less and ended in a dead-end node then it was considered a side-chain and removed from the graph.

### I. Helix Refinement

In the final post-processing step, we tackled prediction inaccuracies for α-helix backbone structures. Due to their geometrical shape, the neural network had, in some cases, difficulties accurately predicting the location of Cα atoms belonging to an α-helix. In order to improve the prediction, we exploited the fact that the shape of an α-helix has a general definition which is valid across proteins [33]. Since the neural network predicted the confidence of secondary structure elements, as described in Network Output section, we know which Cα atoms belong to an α-helix based on the confidence of their region in space. We combined this knowledge of α-helix locations and their shape attributes in order to adjust the appropriate Cα atoms to better fit the shape of a natural α-helix structure.

For an α-helix which centers around the z-axis, we can use Equation 1 to model its shape where the variables *s* and *r* represent the initial shift and rotation of the helix. The values *2.11* and *1.149* are constants that define the radius and pitch of the helix to best match those of an α-helix.

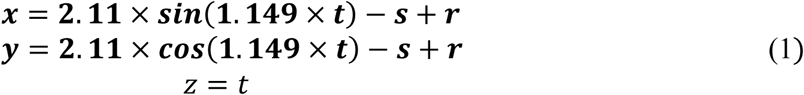

Equation 1 however, cannot be used to describe an α-helix which does not center around the z-axis or whose shape is not a straight cylinder. Since this is the case for most α-helices, it is necessary to adjust the equation in such way that it will address these issues. With the aim of doing so, we first locate the screw axis, the center line around which the helix winds itself, for each α-helix. This is achieved by calculating the centroid of consecutive intervals of the α-helix and then connecting them to approximate the true curve. An example of an α-helix and its calculated screw axis can be seen in Fig. 7.

**Fig. 7.**
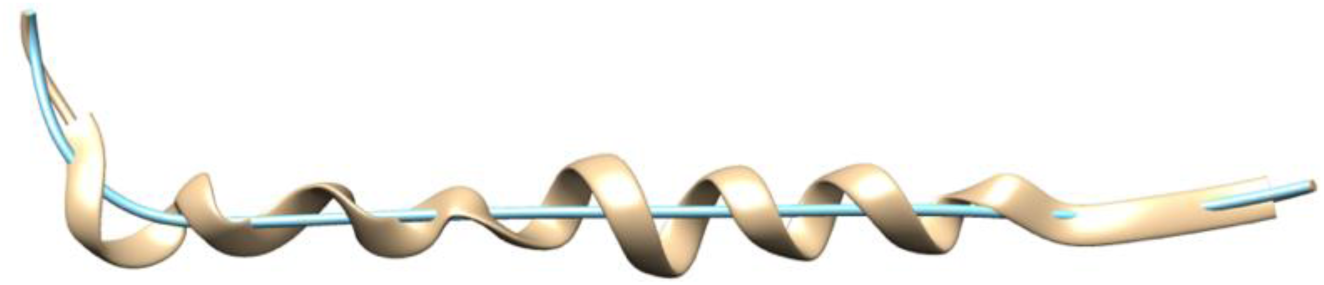
Alpha-Helix extracted from the backbone prediction of the 5u70 protein map in tan color and its screw axis in teal color.

Now that we know the location and shape of the screw axis for the α-helix, we need to incorporate this information into Equation 1. This is achieved by interpreting *t* as the distance that we travelled on the screw axis and use the unit direction vector of the screw axis at a certain point *t* as the new z axis. Next, we can find the new y-axis by calculating the cross product of the x-axis and the new z-axis and then normalizing it. Finally, we can get the new x-axis by calculating the cross product of the new z and y-axis and normalizing it again. By concatenating the three new axes we can get a rotation matrix **RM** with which we can calculate the point of the α-helix for any value *t* as shown in Equation 2.

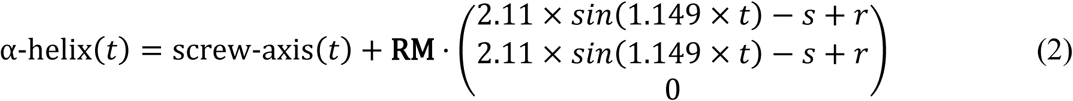

Now, we need to know the values *t* at which we have to insert Cα atoms. Since we know that an α-helix has a rise of 1.5Å per residue [33] we can increase *t* in steps of 1.5 and add a new Cα atom at α-helix(*t*).

In the final step we minimize the average distance from the Cα atoms of the refined α-helix to the Cα atoms of the original prediction. This is done by applying a minimization algorithm^6^ over the variables *s* and *r* to try different initial shifts and rotations. The final results of the α-helix refinement step are shown in Fig. 8.

**Fig. 8.**
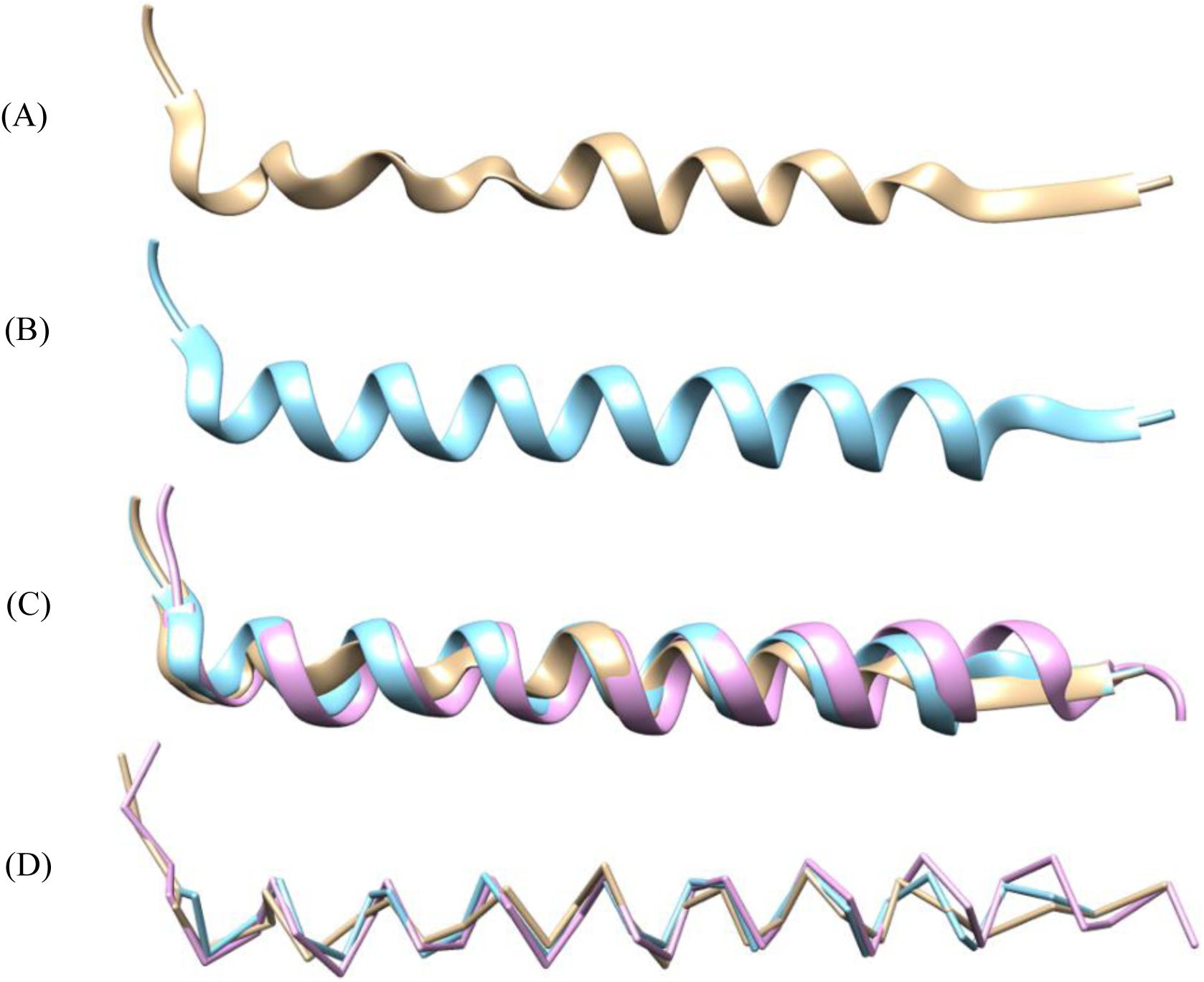
Alpha-Helix extracted from the backbone prediction of the 5u70 protein map. (A) Original prediction before the helix-refinement step. (B) Alpha-Helix after the refinement. (C) Direct comparison of original prediction in tan color, refined prediction colored in teal, and the ground truth in pink color. (D) Direct comparison without ribbon.

### J. Mapping Protein Sequences onto Cα traces

After an imperfectly reconstructed Cα trace is reconstructed, the next important problem is to assign amino acids in the protein sequence onto its correct location in the trace. This problem is non-trivial because a protein may have extra or disordered residues that do not have corresponding positions in the trace. Additionally, the trace contains noisy, false or missing Cα positions that do not match with residues in the protein well. Therefore, simply copying a protein sequence into a Cα trace does not work. To address this challenge, we design a quality assessment-based combinatorial algorithm to map a protein sequence onto each reconstructed Cα trace from the previous steps.

As shown in Fig. 9, given the Cα trace of a target protein and its whole sequence, the algorithm first extracts all continuous Cα segments with length greater than a threshold (i.e. >50 residues). For each Ca segment, a search on the whole sequence is performed to identify its best-matching sequence fragment (sub-sequence) according to the fitness between the Ca segment and the sequence fragment (i.e. the energy or the structural quality score). Specifically, the target protein sequence is decomposed into all possible sequence fragments with the same length of the Cα trace segment and each sequence fragment is mapped into the trace segment. Based on the Cα coordinate of each sequence fragment obtained from the mapping, the main chain structure including the positions of N, Cα, C atoms for the sequence fragment is reconstructed by using Pulchra [34]. Scwrl [35] is then used to add side chains into the main-chain structure of the sequence fragment to obtain a full-atom structure. The quality of the structure of each sequence fragment, i.e. the fitness between the Cα trace and the sequence fragment, is assessed by a protein single-model quality assessment method Qprob [36], which utilizes several structural and physicochemical features by feature-based probability density functions to predict the structure quality score (GDT-TS). Finally, the sequence fragment whose assigned structure has the best structural quality is selected to map to the Cα trace. All the Ca-trace segments were evaluated one by one according to the segment size from largest to smallest to identify their best-matching sequence fragments. If the optimal sequence fragment for current Ca-trace segment has already been assigned to one of previous segments, the unassigned sequence fragment with the largest quality score was then selected for the segment.

**Fig. 9.**
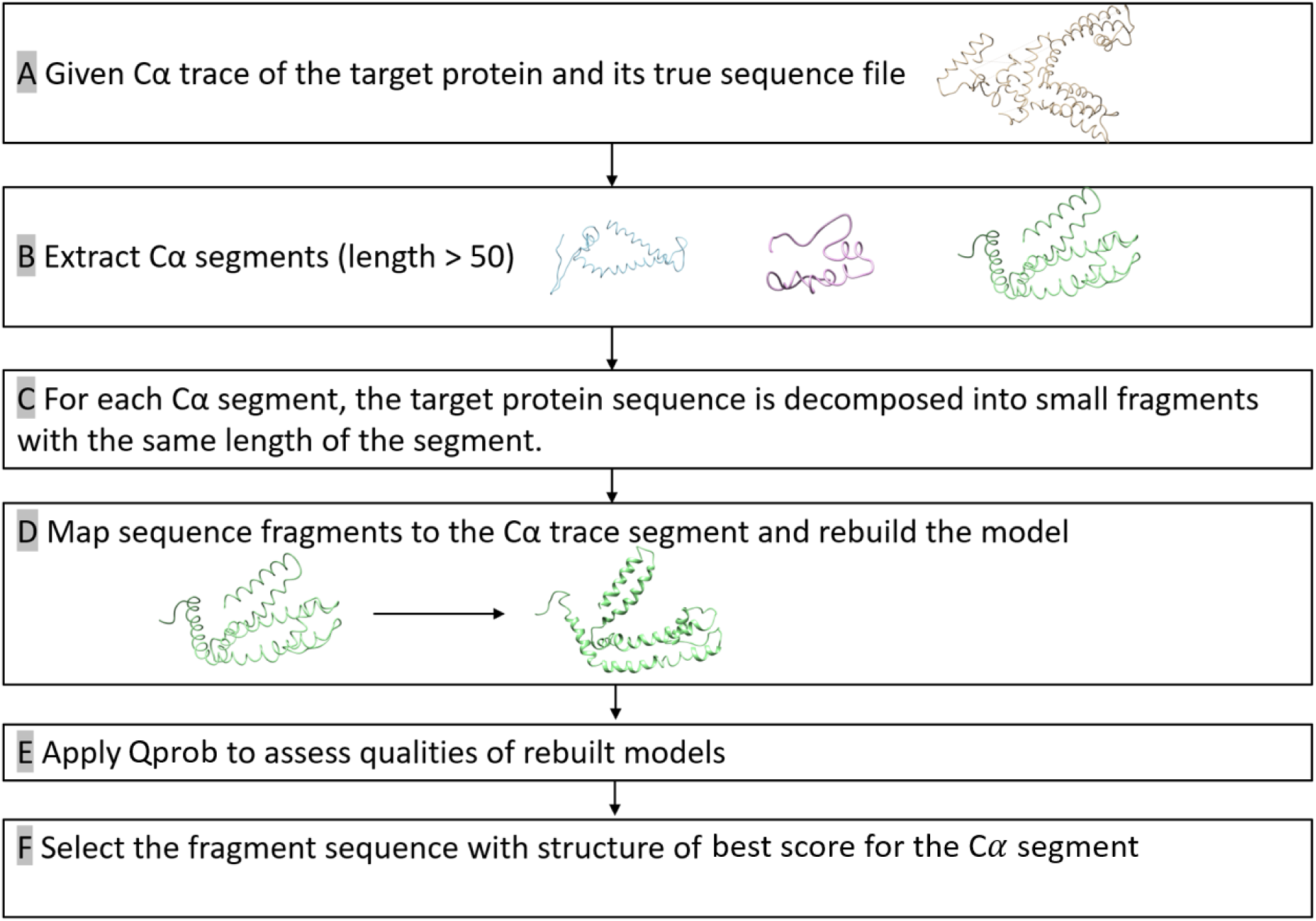
The algorithm of mapping a protein sequence onto a Ca trace.

### K. Computation

The C-CNN was trained with 25,000 simulated protein maps, each with a size of 64×64×64 voxels. Training was accomplished with the Python TensorFlow Library on a Nvidia GTX 1070 GPU. Training was stopped after 15 epochs to prevent overfitting and took about 24 hours. Density Map prediction, which involved running a preprocessed density map through the saved C-CNN only, was completed using the same GPU and took about 15 seconds to produce the five output prediction maps (three SSEs, backbone, and Cα atoms) for a map size of approximately 100×100×100 voxels. The path-walking algorithm was the most time-consuming aspect of the prediction process. A map of approximately 1000 Cα atoms took about 20 minutes to compute. All computation used a machine with an Intel 6 core i7-8700K CPU clocked at 3.7GHz with 16GB of RAM. To increase throughput, a new process was spawned for each path-walking task (one per protein, with a maximum of 12 at a time).

## III. Results

This method was tested with both simulated and experimental maps. Simulated density maps were generated using the *pdb2mrc* script from the EMAN2 package while experimental maps were downloaded from the EM Databank. Both experimental and simulated maps underwent the same pre-processing steps as outlined in the Data Collection/Generation section before being evaluated.

### A. Metrics

A variety of metrics were used to measure the effectiveness of our method. One primary metric was the root-mean-squared-deviation (RMSD) which measures the standard deviation of the distance between atoms in two models. In this case, the two models were the predicted Cα atoms and the ground truth Cα atoms as defined in the PDB file. The output from our model often consisted of partial backbone traces when the confidence was not high enough to form a complete backbone trace. With partial backbone traces it is difficult to use traditional RMSD algorithms to measure the effectiveness of a prediction method. As a result, we decided to follow the same method used by the fully autonomous Phenix method [19] which compares each Cα atom in the ground truth model to the closest Cα atom in the predicted method using a one-to-one mapping. This RMSD method walks each predicted backbone trace and pairs it with the closest Cα atom in the ground truth structure. This produces lower/better RMSD values than other methods [12] [25] because it allows for Cα skips in the ground truth backbone trace.

Another primary metric that we focus on in this research is the percentage of predicted Cα atoms within 3Å (% Cα in 3Å) of the ground truth structure. This metric is a good measure of prediction map completeness because higher values mean that a higher percentage of the ground truth atoms were found. This metric is calculated in a similar way to the RMSD metric: by walking down each predicted trace and pairing each predicted Cα atom with its closest ground truth Cα atom. This metric requires a one-to-one mapping. We compare our results for this metric to the Phenix method which used the exact same metric and metric calculation method.

In addition to these metrics, we also include for each tested density map: the number of predicted Cα atoms, the number of actual Cα atoms, and the number of false positive Cα atoms.

### B. Simulated Density Maps

Table 1 shows the results for seven simulated density maps. Each density map was generated from the *pdb2mrc* script at 6Å. Although 6Å is considered medium resolution, *pdb2mrc* used inflated resolution values. Therefore, a 6Å resolution simulated map was most similar to an experimental map at 3.5Å resolution.

**Table 1.** Results of simulated data using the script pdb2mrc.py from the EMAN2 package at 6Å resolution.

| PDB ID | Model Cα | Native Cα | RMSD (Å) | % Cα in 3Å | FP% |
| --- | --- | --- | --- | --- | --- |
| 3i2n | 351 | 345 | 0.91 | 99.1 | 0.6 |
| 3n2t | 328 | 327 | 0.90 | 99.4 | 0.3 |
| 3qc7 | 165 | 164 | 0.97 | 94.8 | 0.0 |
| 5i68 | 661 | 662 | 0.92 | 99.4 | 0.2 |
| 6ahv | 348 | 345 | 0.89 | 99.7 | 0.3 |
| 6eyw | 384 | 381 | 0.87 | 100.0 | 0.3 |
| 6g6l | 112 | 111 | 0.84 | 98.2 | 0.9 |
| **Avg.** |  |  | **0.90** | **98.7** |  |

The simulated results proved to be very accurate relative to the true protein structure. With an average RMSD of 0.90Å per Cα atom, the predicted backbone structure was almost a perfect match to the true structure. Additionally, the predicted results produced nearly complete backbone traces as evident by an average of 98.7% of ground truth Cα atoms being within 3Å of a predicted Cα atom. Finally, the false positive rate was very low for each prediction (no more than 2 false positive atoms per protein). This translates to a very high sensitivity for each prediction. Fig. 10 shows the final prediction maps of both the Cα-only backbone structure along with the ribbon representation of 6eyw.

**Fig. 10.**
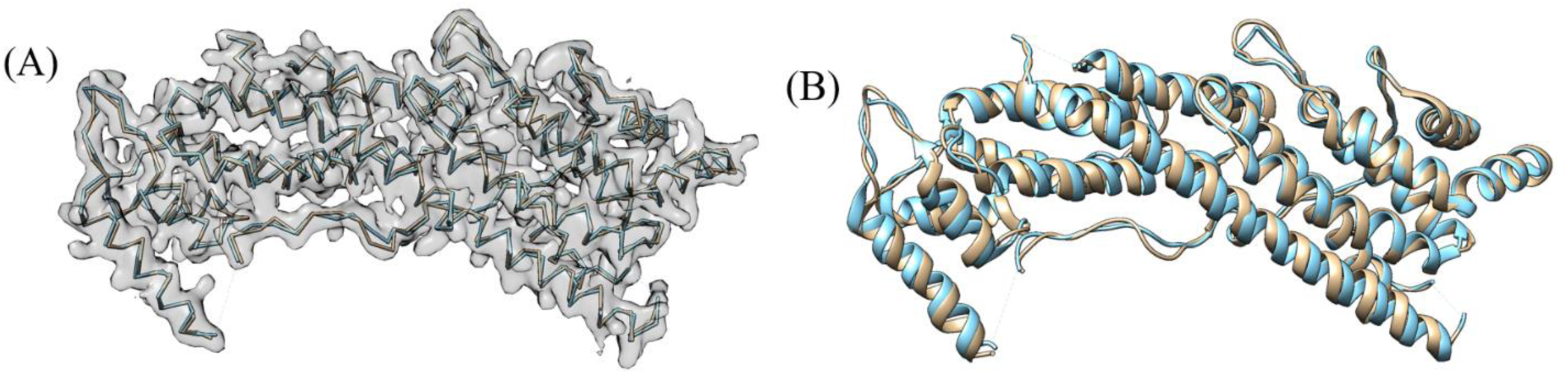
Final Backbone Prediction structures from the simulated 6eyw protein map. Ground Truth is colored teal while the predicted structures are colored tan. (A) contains the Cα only backbone structures with the input density map overlaid on the image for reference. (B) displays the final ribbon prediction. The ribbon was enhanced using the secondary structure output maps from the Cascaded Convolutional Neural Network.

The low RMSD values, minimal false positive Cα atoms, and high matching percentage of Cα atoms within 3Å of the real structure demonstrates the effectiveness of this method with simulated density maps. However, these maps have the advantage over experimental maps in that they do not contain any experimental inaccuracies or real-world noise to distort the image. Additionally, these simulated maps were generated with the same *pdb2mrc* script that the training data was generated with. This means that the C-CNN likely learned features of simulated maps very accurately and possibly overfit the data thereby leading to the high accuracy metrics.

### C. Experimental Density Maps

The real test of our backbone prediction model involved experimental density maps. Experimental maps ranging in resolution from 2.6Å to 4.4Å were evaluated using the backbone prediction model. Table 2 tabulates the results for each experimental density map. Each density map was downloaded from the EM databank^7^. In addition to the normal preprocessing steps outlined in the Pre-Processing section, each density map underwent an additional cleaning step before being evaluated. To remove artifacts from the cyro-EM imaging process unrelated to the protein structure, all voxels that were further than approximately 5Å from the ground truth structure were zeroed out using built in Chimera functions. This cleaning step essentially cut the protein out from the raw density map. However, this step maintained the experimental protein’s shape and density and did not change or manipulate any voxel data within the protein’s experimental structure.

**Table 2.**
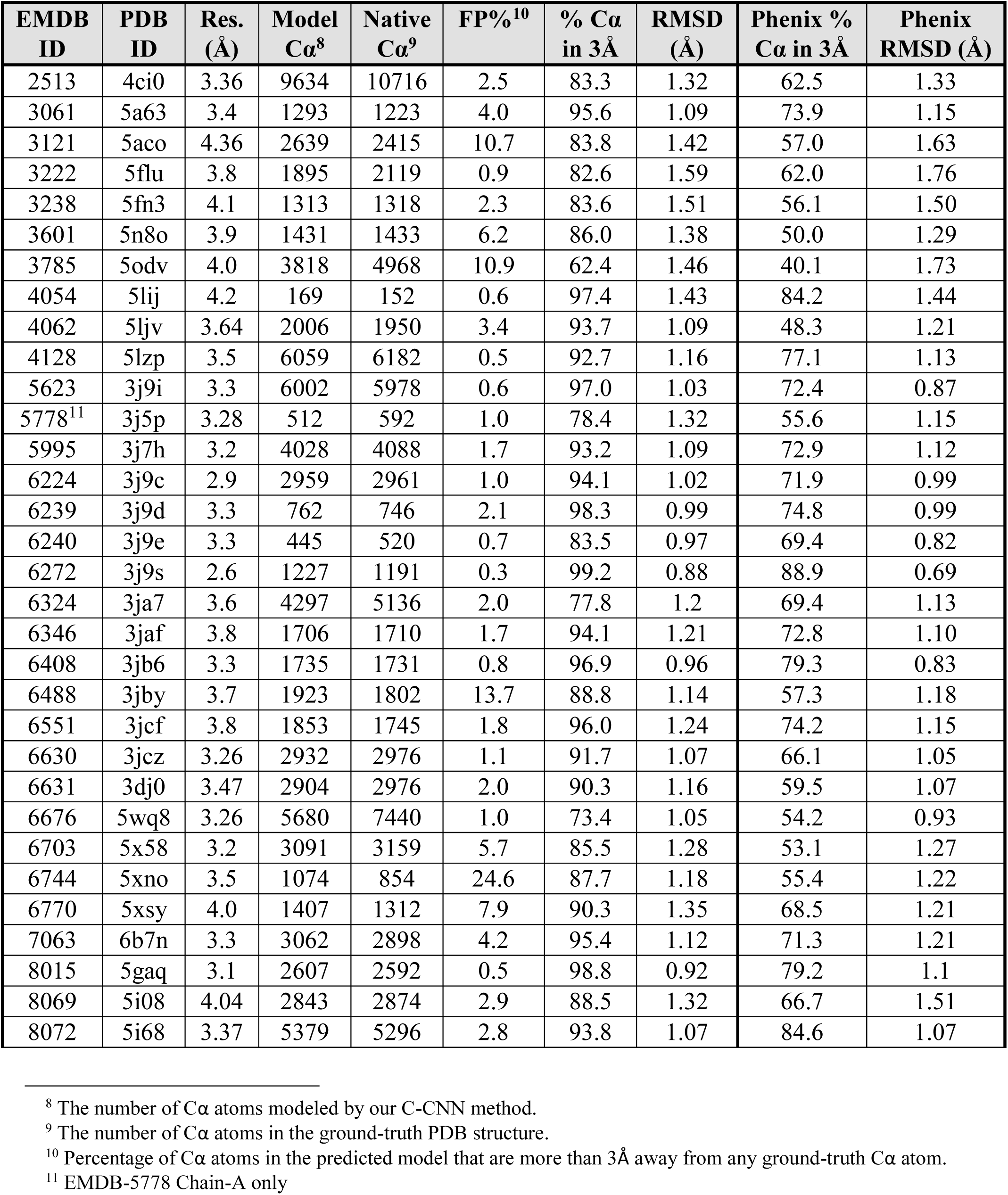
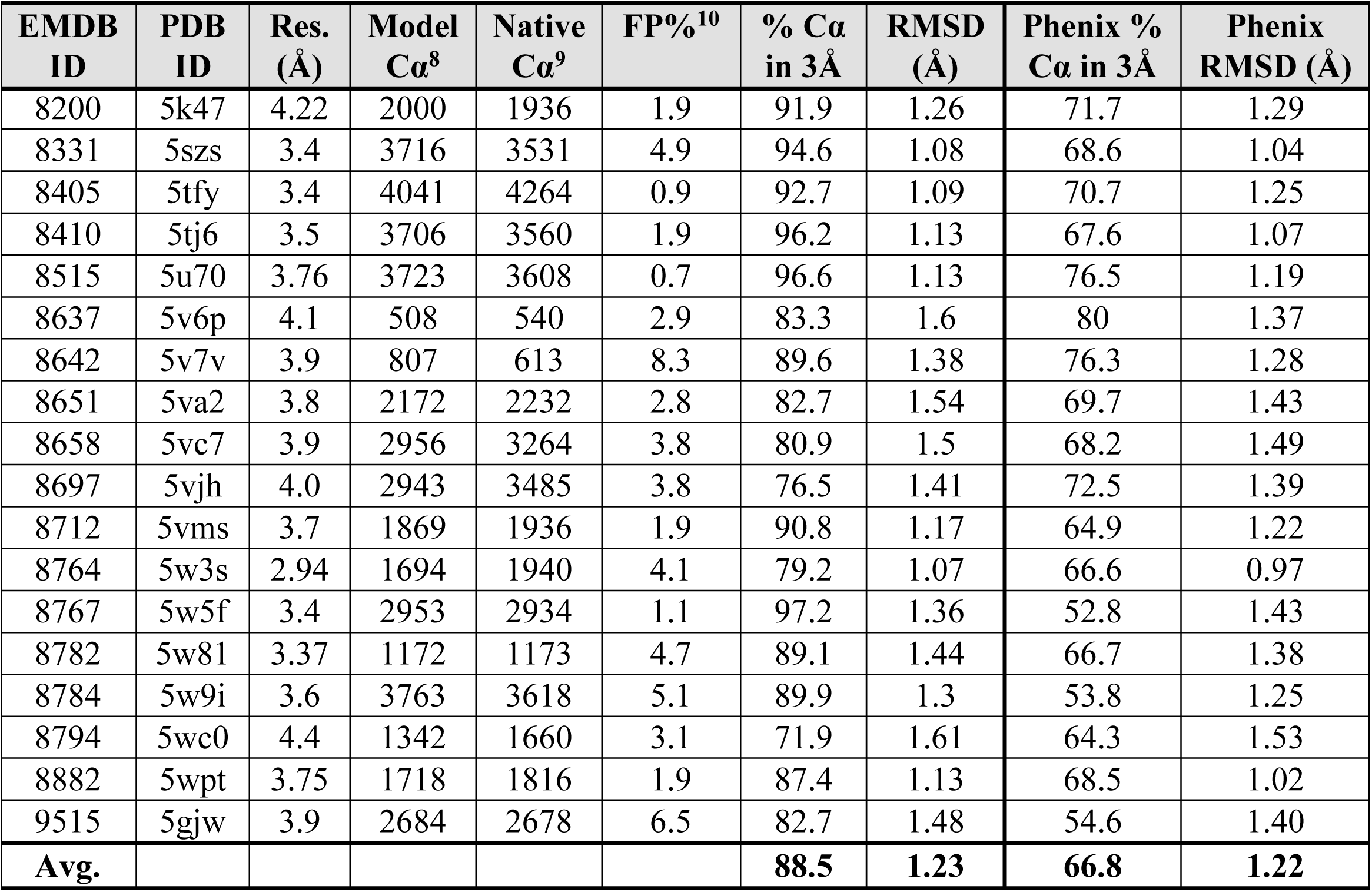
Results from the evaluation of experimental density maps. Results from the Phenix method [19] are listed alongside each evaluated map to compare the RMSD and % matching metrics of each method. Combined metrics for each method are plotted in Fig. 14 and Fig. 15. (Results are continued on the next page)

| EMDB ID | PDB ID | Res. (Å) | Model Cα<sup>8</sup> | Native Cα<sup>9</sup> | FP%<sup>10</sup> | % Cα in 3Å | RMSD (Å) | Phenix % Cα in 3Å | Phenix RMSD (Å) |
| --- | --- | --- | --- | --- | --- | --- | --- | --- | --- |
| 2513 | 4ci0 | 3.36 | 9634 | 10716 | 2.5 | 83.3 | 1.32 | 62.5 | 1.33 |
| 3061 | 5a63 | 3.4 | 1293 | 1223 | 4.0 | 95.6 | 1.09 | 73.9 | 1.15 |
| 3121 | 5aco | 4.36 | 2639 | 2415 | 10.7 | 83.8 | 1.42 | 57.0 | 1.63 |
| 3222 | 5flu | 3.8 | 1895 | 2119 | 0.9 | 82.6 | 1.59 | 62.0 | 1.76 |
| 3238 | 5fn3 | 4.1 | 1313 | 1318 | 2.3 | 83.6 | 1.51 | 56.1 | 1.50 |
| 3601 | 5n8o | 3.9 | 1431 | 1433 | 6.2 | 86.0 | 1.38 | 50.0 | 1.29 |
| 3785 | 5odv | 4.0 | 3818 | 4968 | 10.9 | 62.4 | 1.46 | 40.1 | 1.73 |
| 4054 | 5lij | 4.2 | 169 | 152 | 0.6 | 97.4 | 1.43 | 84.2 | 1.44 |
| 4062 | 5ljv | 3.64 | 2006 | 1950 | 3.4 | 93.7 | 1.09 | 48.3 | 1.21 |
| 4128 | 5lzp | 3.5 | 6059 | 6182 | 0.5 | 92.7 | 1.16 | 77.1 | 1.13 |
| 5623 | 3j9i | 3.3 | 6002 | 5978 | 0.6 | 97.0 | 1.03 | 72.4 | 0.87 |
| 5778<sup>11</sup> | 3j5p | 3.28 | 512 | 592 | 1.0 | 78.4 | 1.32 | 55.6 | 1.15 |
| 5995 | 3j7h | 3.2 | 4028 | 4088 | 1.7 | 93.2 | 1.09 | 72.9 | 1.12 |
| 6224 | 3j9c | 2.9 | 2959 | 2961 | 1.0 | 94.1 | 1.02 | 71.9 | 0.99 |
| 6239 | 3j9d | 3.3 | 762 | 746 | 2.1 | 98.3 | 0.99 | 74.8 | 0.99 |
| 6240 | 3j9e | 3.3 | 445 | 520 | 0.7 | 83.5 | 0.97 | 69.4 | 0.82 |
| 6272 | 3j9s | 2.6 | 1227 | 1191 | 0.3 | 99.2 | 0.88 | 88.9 | 0.69 |
| 6324 | 3ja7 | 3.6 | 4297 | 5136 | 2.0 | 77.8 | 1.2 | 69.4 | 1.13 |
| 6346 | 3jaf | 3.8 | 1706 | 1710 | 1.7 | 94.1 | 1.21 | 72.8 | 1.10 |
| 6408 | 3jb6 | 3.3 | 1735 | 1731 | 0.8 | 96.9 | 0.96 | 79.3 | 0.83 |
| 6488 | 3jby | 3.7 | 1923 | 1802 | 13.7 | 88.8 | 1.14 | 57.3 | 1.18 |
| 6551 | 3jcf | 3.8 | 1853 | 1745 | 1.8 | 96.0 | 1.24 | 74.2 | 1.15 |
| 6630 | 3jcz | 3.26 | 2932 | 2976 | 1.1 | 91.7 | 1.07 | 66.1 | 1.05 |
| 6631 | 3dj0 | 3.47 | 2904 | 2976 | 2.0 | 90.3 | 1.16 | 59.5 | 1.07 |
| 6676 | 5wq8 | 3.26 | 5680 | 7440 | 1.0 | 73.4 | 1.05 | 54.2 | 0.93 |
| 6703 | 5x58 | 3.2 | 3091 | 3159 | 5.7 | 85.5 | 1.28 | 53.1 | 1.27 |
| 6744 | 5xno | 3.5 | 1074 | 854 | 24.6 | 87.7 | 1.18 | 55.4 | 1.22 |
| 6770 | 5xsy | 4.0 | 1407 | 1312 | 7.9 | 90.3 | 1.35 | 68.5 | 1.21 |
| 7063 | 6b7n | 3.3 | 3062 | 2898 | 4.2 | 95.4 | 1.12 | 71.3 | 1.21 |
| 8015 | 5gaq | 3.1 | 2607 | 2592 | 0.5 | 98.8 | 0.92 | 79.2 | 1.1 |
| 8069 | 5i08 | 4.04 | 2843 | 2874 | 2.9 | 88.5 | 1.32 | 66.7 | 1.51 |
| 8072 | 5i68 | 3.37 | 5379 | 5296 | 2.8 | 93.8 | 1.07 | 84.6 | 1.07 |

<sup>8</sup> The number of Cα atoms modeled by our C-CNN method.
<sup>9</sup> The number of Cα atoms in the ground-truth PDB structure.
<sup>10</sup> Percentage of Cα atoms in the predicted model that are more than 3Å away from any ground-truth Cα atom.
<sup>11</sup> EMDB-5778 Chain-A only

| EMDB ID | PDB ID | Res. (Å) | Model Cα<sup>8</sup> | Native Cα<sup>9</sup> | FP%<sup>10</sup> | % Cα in 3Å | RMSD (Å) | Phenix % Cα in 3Å | Phenix RMSD (Å) |
| --- | --- | --- | --- | --- | --- | --- | --- | --- | --- |
| 8200 | 5k47 | 4.22 | 2000 | 1936 | 1.9 | 91.9 | 1.26 | 71.7 | 1.29 |
| 8331 | 5szs | 3.4 | 3716 | 3531 | 4.9 | 94.6 | 1.08 | 68.6 | 1.04 |
| 8405 | 5tfy | 3.4 | 4041 | 4264 | 0.9 | 92.7 | 1.09 | 70.7 | 1.25 |
| 8410 | 5tj6 | 3.5 | 3706 | 3560 | 1.9 | 96.2 | 1.13 | 67.6 | 1.07 |
| 8515 | 5u70 | 3.76 | 3723 | 3608 | 0.7 | 96.6 | 1.13 | 76.5 | 1.19 |
| 8637 | 5v6p | 4.1 | 508 | 540 | 2.9 | 83.3 | 1.6 | 80 | 1.37 |
| 8642 | 5v7v | 3.9 | 807 | 613 | 8.3 | 89.6 | 1.38 | 76.3 | 1.28 |
| 8651 | 5va2 | 3.8 | 2172 | 2232 | 2.8 | 82.7 | 1.54 | 69.7 | 1.43 |
| 8658 | 5vc7 | 3.9 | 2956 | 3264 | 3.8 | 80.9 | 1.5 | 68.2 | 1.49 |
| 8697 | 5vjh | 4.0 | 2943 | 3485 | 3.8 | 76.5 | 1.41 | 72.5 | 1.39 |
| 8712 | 5vms | 3.7 | 1869 | 1936 | 1.9 | 90.8 | 1.17 | 64.9 | 1.22 |
| 8764 | 5w3s | 2.94 | 1694 | 1940 | 4.1 | 79.2 | 1.07 | 66.6 | 0.97 |
| 8767 | 5w5f | 3.4 | 2953 | 2934 | 1.1 | 97.2 | 1.36 | 52.8 | 1.43 |
| 8782 | 5w81 | 3.37 | 1172 | 1173 | 4.7 | 89.1 | 1.44 | 66.7 | 1.38 |
| 8784 | 5w9i | 3.6 | 3763 | 3618 | 5.1 | 89.9 | 1.3 | 53.8 | 1.25 |
| 8794 | 5wc0 | 4.4 | 1342 | 1660 | 3.1 | 71.9 | 1.61 | 64.3 | 1.53 |
| 8882 | 5wpt | 3.75 | 1718 | 1816 | 1.9 | 87.4 | 1.13 | 68.5 | 1.02 |
| 9515 | 5gjw | 3.9 | 2684 | 2678 | 6.5 | 82.7 | 1.48 | 54.6 | 1.40 |
| **Avg.** |  |  |  |  |  | **88.5** | **1.23** | **66.8** | **1.22** |

Our results were compared to the fully automatic Phenix method [19] in two categories: RMSD and % Cα Matching within 3Å. The Phenix method results are listed alongside our results in Table 2.

The results on experimental data show that our method was very similar to the automatic Phenix method with respect to RMSD. The C-CNN was able to achieve a mean RMSD of 1.23Å while the Phenix method achieve a mean RMSD of 1.22Å. Both methods were tested on the same set of 50 experimental density maps. Our method was able to produce a much higher average Cα percentage matching within 3Å than the Phenix method (88.5% vs. 66.8%). This significant improvement is primarily a result of our method’s ability to predict backbone structure in relatively low-confident regions while the automatic Phenix made no prediction in these areas. By achieving similar RMSD metrics but improved Cα matching percentages our method has clearly demonstrated an improvement over the automatic Phenix method in terms of Cα/backbone prediction. Fig. 11 shows the final predicted map for three experimental density maps using our deep learning method.

**Fig. 11.**
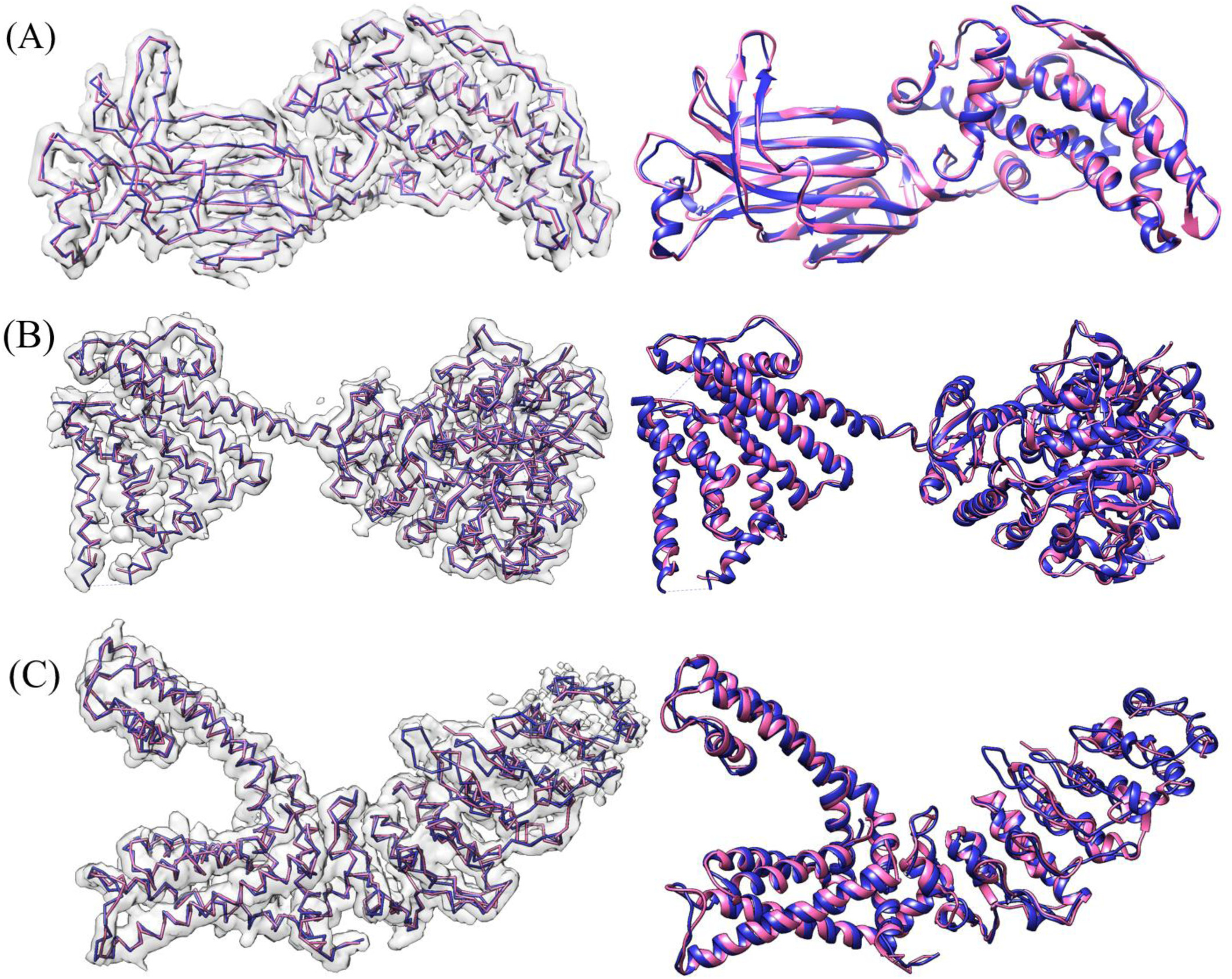
Final Backbone Prediction maps for various density maps. (A) EMDB-6272 (chain-A) at resolution 2.6Å. (B) EMDB-8410 (chain-A) at resolution 3.5Å (C) EMDB-5778 (chain-A), resolution 3.3Å. The left map in each subfigure contains the predicted vs. actual backbone structure. The right map in each subfigure contains the predicted vs. actual ribbon structure of the protein which specify the SSE classification. The pick trace is the predicted structures while the blue trace is the actual structure.

### D. Impact of Helix Refinement

In the Helix Refinement section we discussed the final post-processing step which was responsible for adjusting predicted α-helices to fit the true structure of an α-helix. The refinement step improved the percentage of Cα atoms predicted within 3Å of their actual location by 1.5% while remaining the average RMSD value. In this section we want to highlight specific protein maps where the helix refinement yielded significantly improved results. In Table 3 we can see a comparison of results before and after the application of the helix refinement.

**Table 3.** Results from the evaluation of experimental density maps before and after the helix refinement was applied

| EMDB ID | RMSD before refinement | RMSD after refinement | % Cα in 3Å before refinement | % Cα in 3Å after refinement |
| --- | --- | --- | --- | --- |
| 8637 | 1.68 | 1.60 | 68.5 | 83.3 |
| 3238 | 1.54 | 1.51 | 80.0 | 83.6 |
| 6744 | 1.24 | 1.18 | 84.8 | 87.7 |
| 8200 | 1.30 | 1.26 | 87.9 | 91.9 |
| 8405 | 1.12 | 1.09 | 89.9 | 92.7 |

From Table 3 we can see that the helix refinement was particularly effective in increasing the percentage of Cα atoms predicted within 3Å of their actual location with up to ∼15% in the case of the EMDB-8637 map. In Fig. 12 we can see a comparison before and after the helix refinement step for this map. Improvements in RMSD values were less significant, especially if we consider all test results. This can be attributed to the number of Cα atoms originally predicted within α-helices which is almost always lower than the actual number of Cα atoms. This means that the average inaccuracy in α-helices, which is usually high, has less of an impact to the overall average RMSD value. In the helix refinement process new Cα atoms are added to the predicted α-helices in an attempt to approximate their true structure. As a result, the average RMSD value within the α-helix improves. However, since there are now more Cα atoms belonging to the α-helices, their average inaccuracy has a larger impact to the overall average RMSD dampening improvements.

**Fig. 12.**
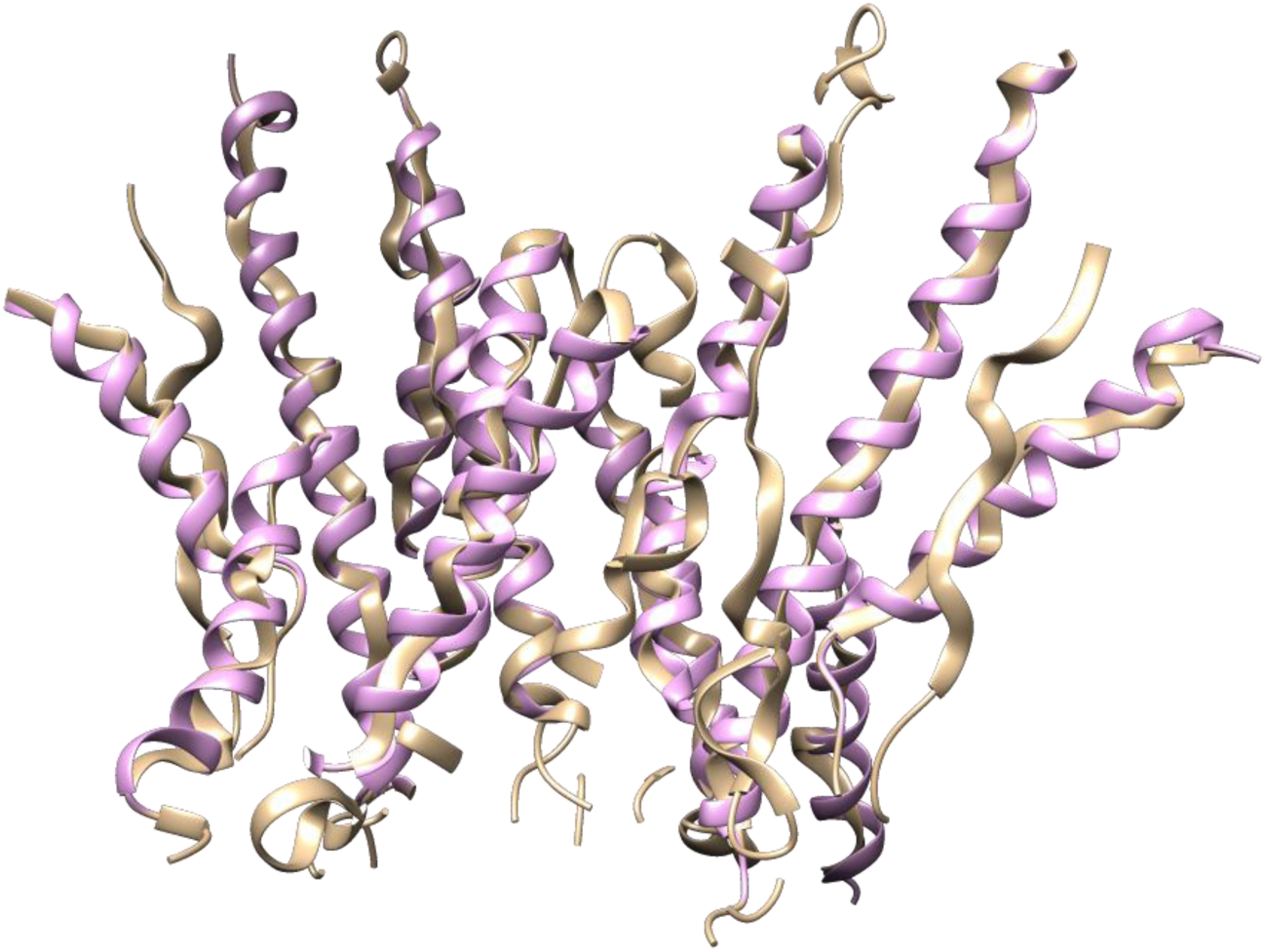
Comparison of predicted map for EMDB-8637 before the helix refinement step in tan color and after the refinement in pink color

### E. Comparison of Prediction Methods

As introduced in the Current Protein Prediction Models section, there are three other leading backbone prediction models: Phenix, MAINMAST, and RosettaES. We compare our deep learning method to the other leading prediction models by evaluating EMDB-5778 (chain-A) with each method. The prediction maps for each method are shown in Fig. 13.

**Fig. 13.**
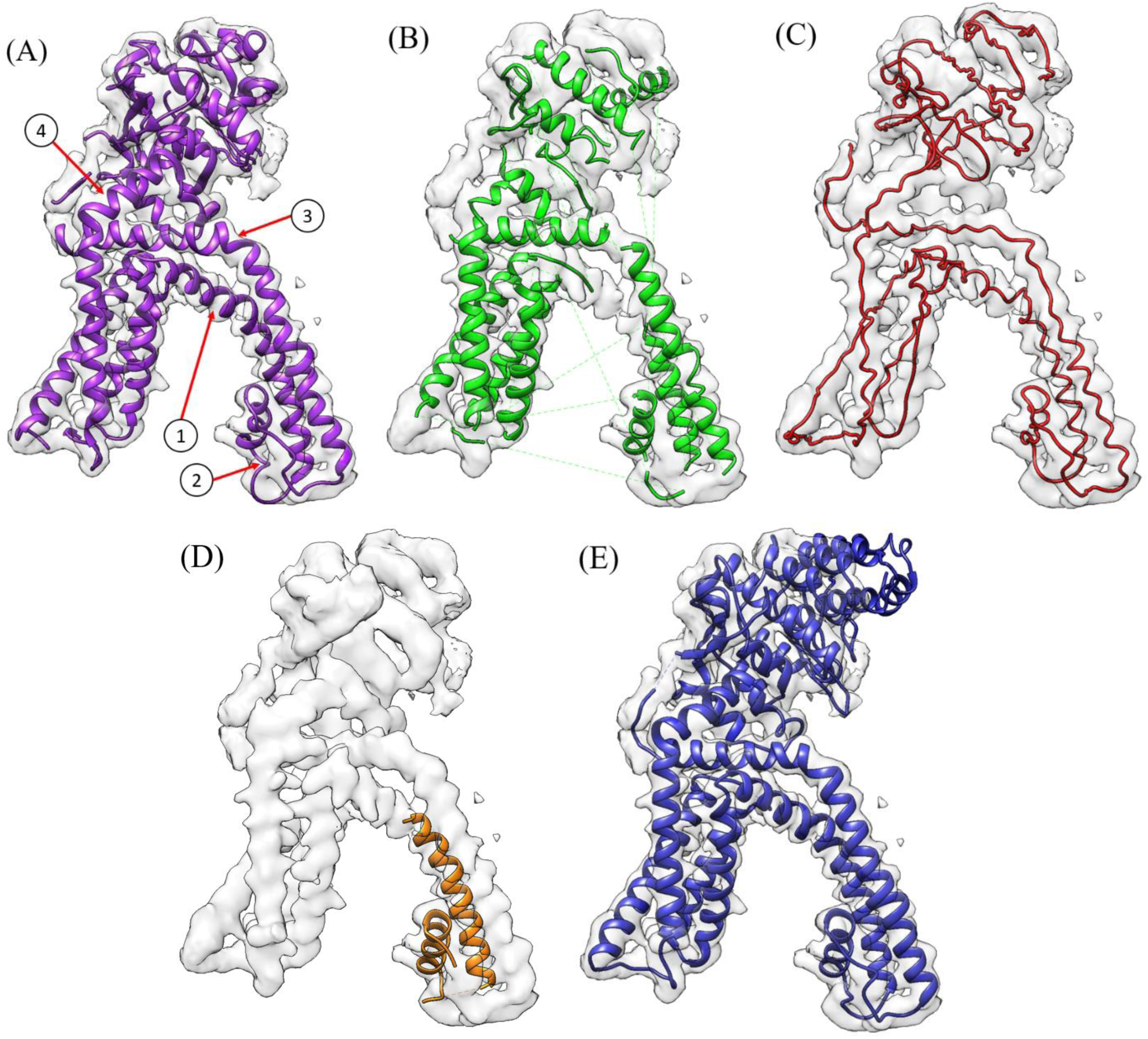
Final Prediction Model for EMDB-5778 (chain-A) at 3.28Å resolution. (A) Our C-CNN method. (B) automatic Phenix method. (C) MAINMAST. (D) RosettaES. (E) The ground truth model. Each figure overlays the input density map on top of the predicted structure. (A) Notes five areas where the C-CNN found the true backbone structure while the automatic Phenix method did not.

Table 4 compares the prediction metrics of all four prediction methods. The RosettaES model produced the most accurate prediction map but it was only able to predict 7.9% of the Cα Atoms from the ground-truth model. The MAINMAST method, which only predicts backbone structure and not SSEs, found the highest percentage of Cα Atoms among the three previous prediction methods. However, MAINMAST has a relatively high RMSD of 2.20Å and also produced the highest false positive Cα Atom percentage. Among the three former prediction methods the Phenix method was arguably the best method for EMDB-5778, producing decent metrics in all three categories (Percentage of Matching Cα Atom, RMSD, and false positive Cα Atom percentage). However, our deep-learning method was about to achieve a much higher Percent Cα Atom Matching and a lower false positive Cα Atom percentage with only a small increase in the RMSD. Fig. 13A points out four areas where our prediction model outperformed the Phenix method by producing a more-complete prediction map.

**Table 4.** Comparison of EMDB-5778 (chain-A) among leading backbone prediction models. Each model had specific strengths, however the deep learning C-CNN produced the most complete model as measured by the percent Cα matching percentage.

| Method | % Cα<br>in 3Å | RMSD<br>(Å) | FP% |
| --- | --- | --- | --- |
| C-CNN | 78.4 | 1.32 | 1.0 |
| Phenix | 55.6 | 1.15 | 3.0 |
| MAINMAST | 59.6 | 2.20 | 6.5 |
| RosettaES | 7.9 | 0.91 | 0.0 |

### F. Computation Time of Prediction Models

While our C-CNN model took about 24 hours to train on a single machine with a GTX 1070 GPU, after the training had completed, the full end-to-end prediction for a density map with approximately 1000 Cα Atoms took about 20 minutes to complete. The C-CNN prediction software was running on the same GTX 1070 GPU with a single CPU core. In contrast, the existing methods usually required a lot of computing resources and were also very time-consuming. In our experiments, RosettaES took 5 days with 20 CPUs to complete, MAINMAST finished in about 18 hours with 1 CPU, and Phenix method took several hours with 6 CPUs.

### G. Comparison of Deep-Learning C-NN and the Fully Automatic Phenix Method

The results for each prediction map in Table 2 are plotted and shown in Fig. 14 and Fig. 15. Fig. 14 plots the RMSD vs. the labeled resolution for each density map. The mean RMSD of our deep-learning method was 1.23Å which is similar to the Phenix method’s RMSD of 1.22Å for the same set of experimental density maps. Fig. 14 shows that the performance of our deep-learning method was very similar to the Phenix method across all resolution values. Fig. 14 also clearly shows how the RMSD for both prediction methods increased as a function of the labeled resolution. This is to be expected as higher resolution maps usually have less well-defined structure to aid the prediction models.

**Fig. 14.**
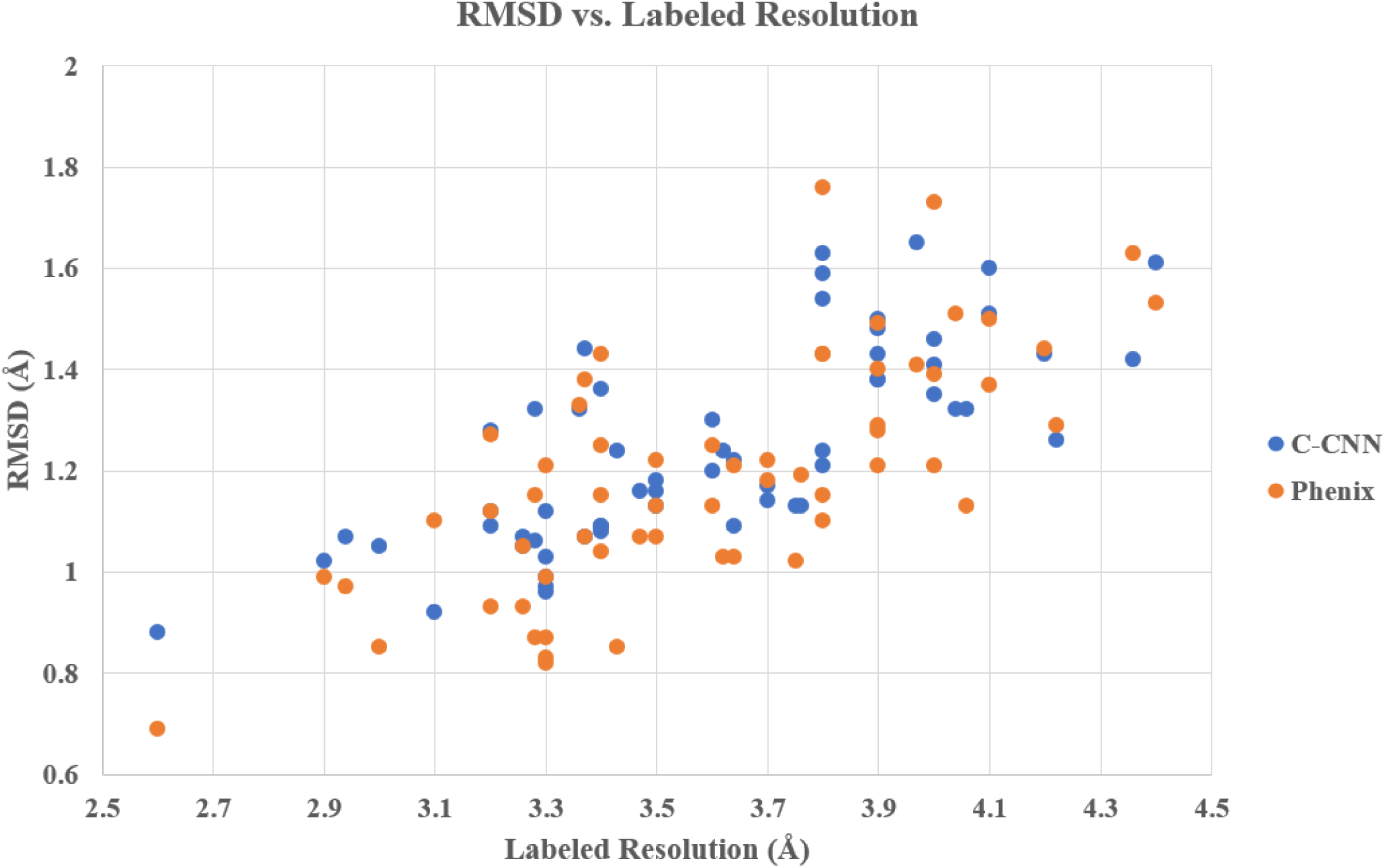
Plot of RMSD as a function of resolution for the 50 experimental density maps. This compares our deep learning C-CNN method and the fully automatic Phenix Method.

**Fig. 15.**
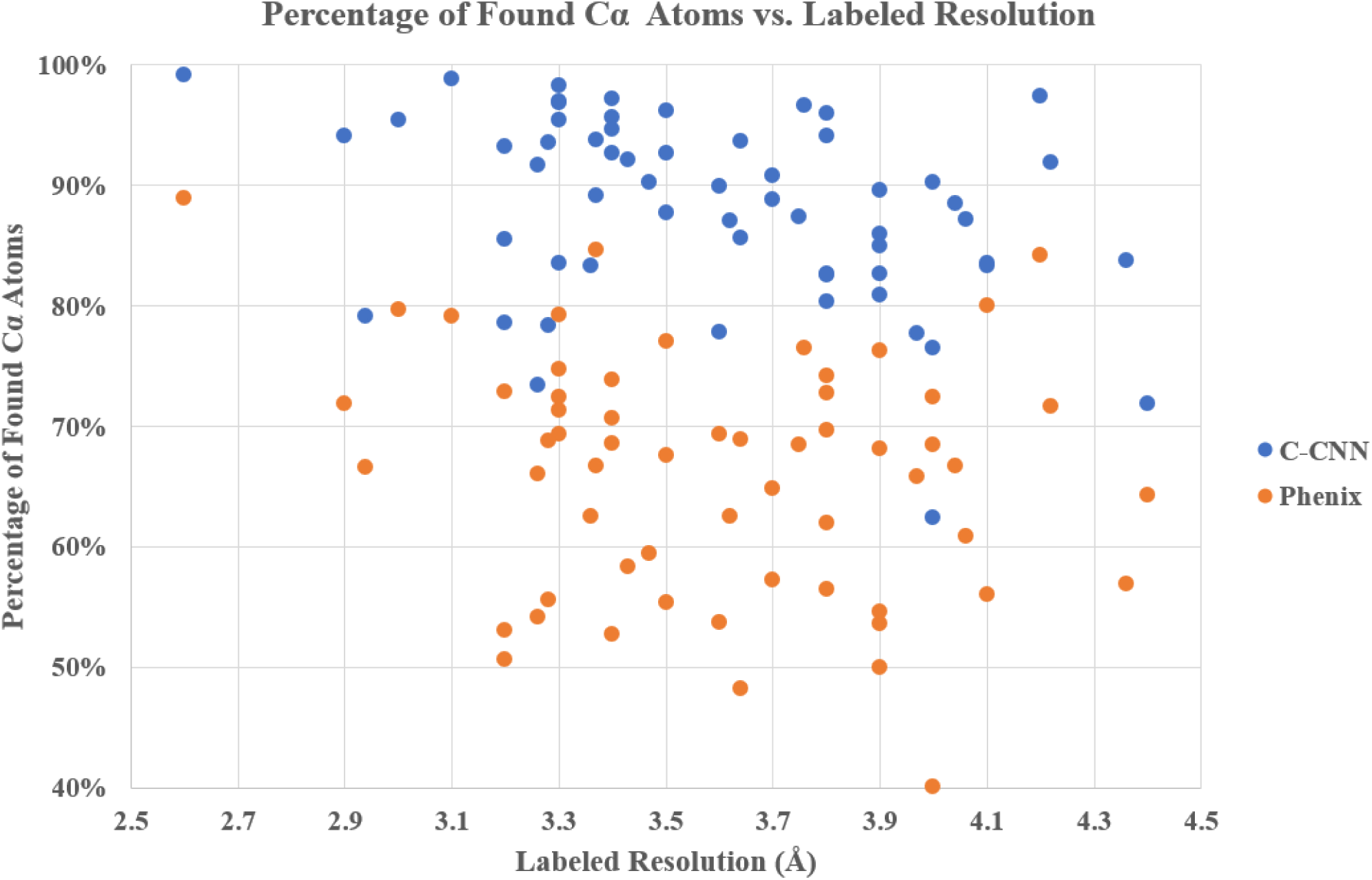
Plot of Percentage of Found Cα atoms as a function of resolution for the 50 experimental density maps. This compares our deep learning C-CNN method and the fully automatic Phenix Method.

Fig. 15 plots the percentage of matching Cα atoms vs. labeled resolution of each density map for both the C-CNN method and the Phenix method. This figure shows how our deep-learning method found a higher percentage of Cα atoms than the Phenix method across all resolutions. Our method found a mean of 88.5% of the Cα atoms for the 50 experimental density maps. This is significantly better that the Phenix method which only found a mean of 66.8% Cα atoms from the same 50 experimental maps.

### H. Evaluating the Results of Mapping Protein Sequences onto Cα traces

To validate the effectiveness of our sequence-to-trace mapping algorithm, we evaluated the structural similarity between the predicted structure of the Cα-trace segment and its real structure for the mapped sequence fragment in the known experimental structure of the protein in terms of the TM-score and GDT-TS scoring metrics. TM-score [37] and GDT-TS [38] scores are two structural similarity measurements with values in (0,1], where higher value indicates better accuracy and 1 means the perfect match between two protein structures. The two metrics measure the match of the residues with Ca-atom distances within a certain distance cutoff of their positions in two structures for the same protein sequence, which is suitable for the method evaluation in our study. A perfect Cα trace segment matched with a completely correct sequence fragment will match perfectly with its corresponding counterpart in the experimental structure, leading to a prefect similarity score (TM-score or GDT-TS score) of 1, otherwise a score between 0 and 1. We validated our mapping algorithm on two targets, EMDB-5778 and EMDB-8410, and the results were summarized in Table 5 and Table 6. For EMDB-5778, the longest Cα trace segment (185 residues) among the three predicted Ca segments is well mapped by the sequence segment and has a TM-score of 0.9478 and a GDT-TS score of 0.8723. The alignment of the mapped structure of the segment with its counterpart, which was superimposed by the sequence-dependent alignment program– TM-score, is illustrated in Fig. 16. The TM-score of the second mapped Cα segment is 0.3947, suggesting its incorrect topology when the TM-score is less than 0.5. For the third mapped Cα segment, the TM-score for the structural alignment is 0.5978, which suggests the close topology is predicted for the sequence fragment. The structural errors result from either the noise in the Cα trace or inaccuracy of the mapping algorithm, or both factors. For EMDB-8410, the longest Cα trace segment (234 residues) is mapped reasonably well, which has a TM-score of 0.8816. However, the other mapped Cα trace segments have TM-scores below the 0.5 threshold, indicating the structures are not predicted correctly. The superposition of the mapped structure and its counterpart for the longest trace of EMDB-8410 is illustrated in Fig. 17. The analysis demonstrates that our mapping strategy is able to identify the correct sequence fragment for the Ca trace if the segment is well predicted by C-CNN and long enough for mapping sequence to structure using quality assessment methods.

**Table 5.** Similarity between three mapped Cα segments (length > 50) and their true counterparts on EMDB-5778

| **EMDB-5778** | **# Ca Atoms** | **TM-score** | **GDT-TS-score** |
| --- | --- | --- | --- |
| Seg #1 | 185 | 0.9478 | 0.8723 |
| Seg #2 | 134 | 0.3947 | 0.3601 |
| Seg #3 | 53 | 0.5978 | 0.6981 |

**Table 6.** Similarity between two mapped Cα segments (length > 50) and their true counterparts on EMDB-8410

| EMDB-8410 | # Ca Atoms | TM-score | GDT-TS-score |
| --- | --- | --- | --- |
| Seg #1 | 234 | 0.8816 | 0.7927 |
| Seg #2 | 180 | 0.4226 | 0.3142 |
| Seg #3 | 159 | 0.4234 | 0.3296 |

**Fig. 16.**
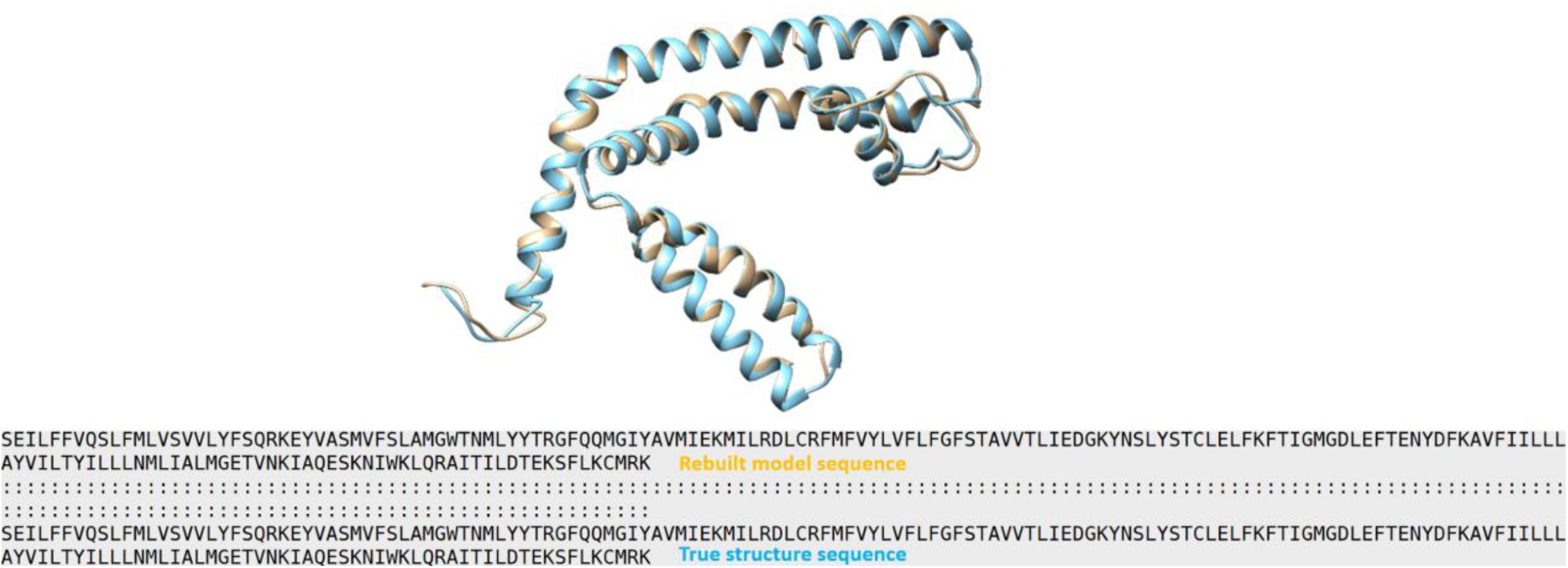
Visualization of superposition between the mapped Cα Segment 1 of EMDB-5778(brown) and its counterpart in the experimental structure (blue). “:” in the sequence-dependent structure alignment denotes the residue pairs whose distance is < 5.0 Angstrom.

**Fig. 17.**
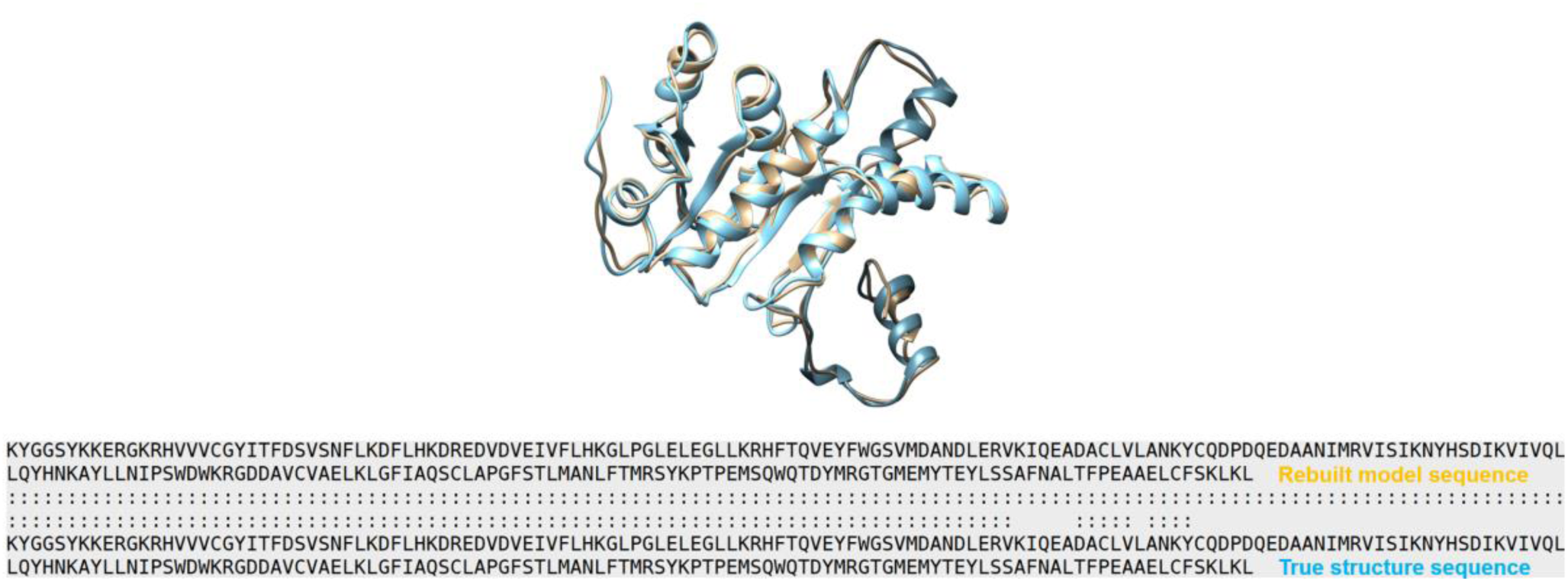
Visualization of the superposition between the mapped Cα segment 1 of EMDB-8410 (brown) and its counterpart experimental structure (blue). “:” denotes the residue pairs of distance < 5.0 Angstrom.

## IV. Discussion

### A. False Positive Cα Error Rate

The RMSD metric in conjunction with the percentage of matching Cα metric is useful to determine the accuracy of Cα atom placements that are in-line with Cα atoms within the ground truth model. However, as is often the case with backbone prediction, side-chains or other noise in a density map will appear as backbone structure to a prediction model. This can cause the model to produce false positive Cα atoms that are clearly not in alignment with the true backbone structure of the protein, see Fig. 18. An effective prediction model will limit the number of false positive Cα atoms by correctly distinguishing side-chains or other noise from the true backbone structure and not place Cα atoms in these areas. In this research, we measure the number of false positive Cα atom for each density map which is listed in Table 1 and Table 2. A false positive is defined as any Cα atom that is placed more than 3Å from any Cα atom within the ground truth structure.

**Fig. 18.**
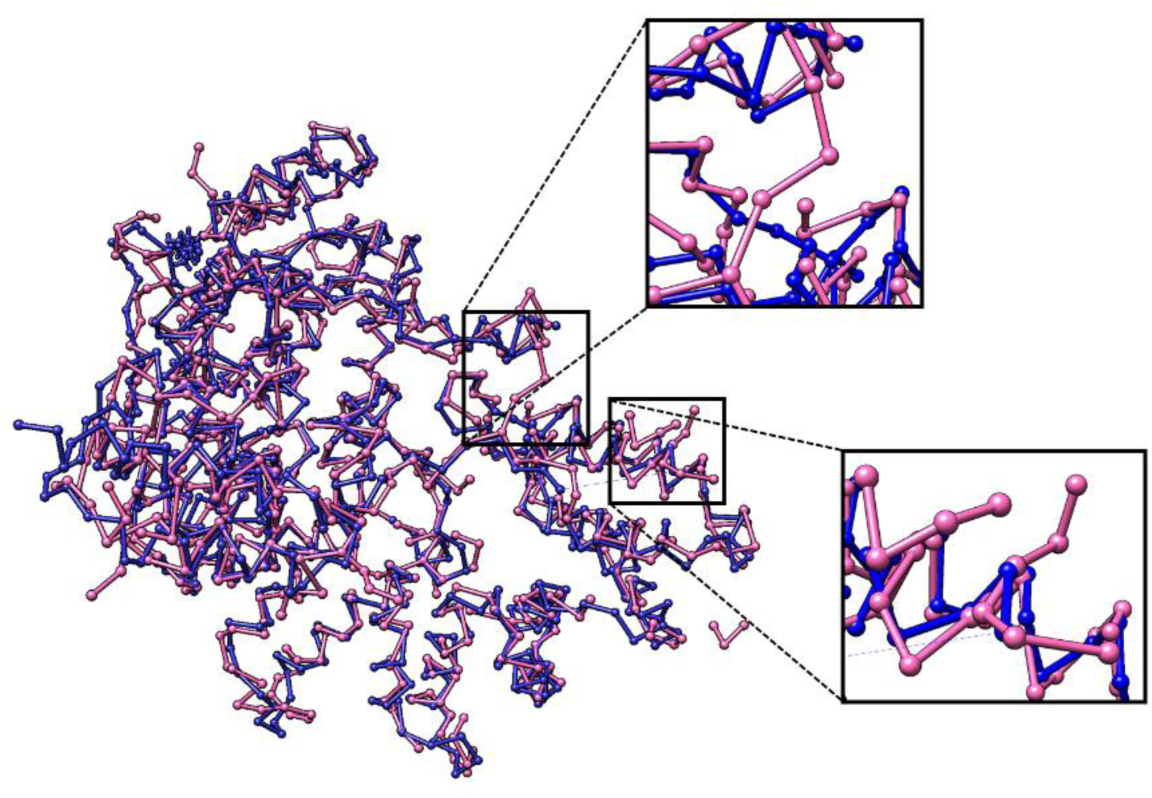
Ball and Stick representation of the prediction map (pink) and the ground truth map (blue) of EMDB-8642, resolution 3.9Å. This map had a relatively low RMSD value of 1.28Å but also had a false positive Cα rate of 8.3%. The expanded areas show examples of side chains that were incorrectly predicted as backbone structure resulting in false-positive Cα placements.

Normalizing the number of false positive Cα atoms by the size of a protein produces a false positive rate for Cα atom placement. This is called the Cα atom error rate. This metric can be derived from the number of false-positive Cα atoms and the total number of predicted Cα atoms, see Equation 3.

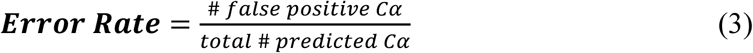

Fig. 19 plots the Cα atom error rate as a function of resolution for the 50 experimental density maps evaluated by our model. The error rate of Cα atom prediction is inversely related to the labeled resolution of a density map. However, our deep learning model produced a relatively low error rate (< ∼5%) for most density maps with a labeled resolution value of 3.7Å or better. The error rate of density maps with a lower resolution than 3.7Å had a high variance and thus had lower prediction confidence.

**Fig. 19.**
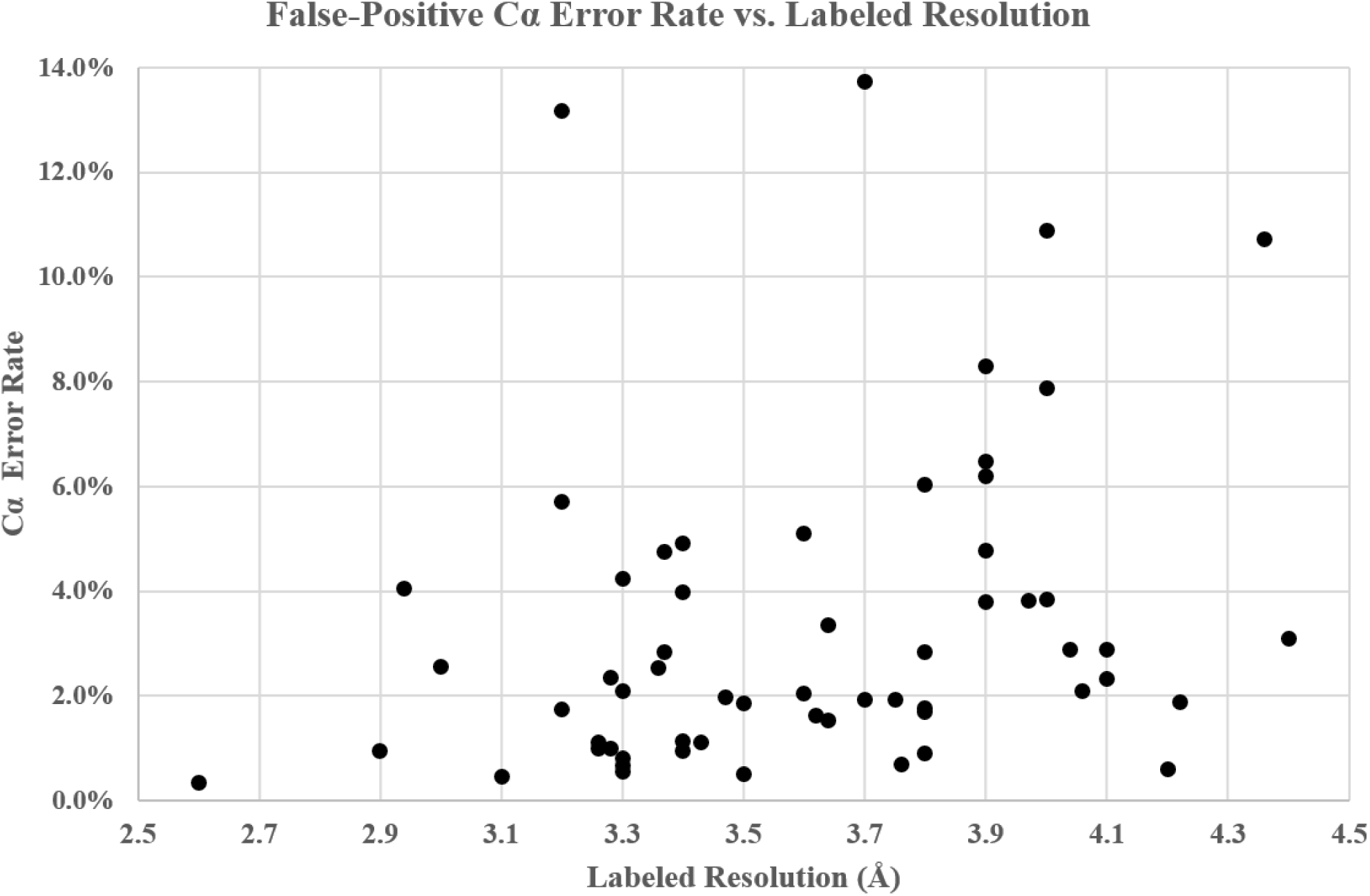
Error-Rate vs. resolution for each experimental prediction map with an added exponential trendline.

#### I. Future Improvements

We found during development that the biggest improvements in accuracy came as the result of adding more convolutional neural networks to the C-CNN. Originally, this method used only one network, the Cα-Atom prediction network. It was only after adding the SSE and Backbone CNNs that we were able to achieve the results outlined in this paper. Future work might be able to incorporate other CNNs into the C-CNN such as an Amino Acid network or an individual atom network. Adding these networks might also help match the sequenced DNA of the protein to the density map.

As the number of publicly available experimental density maps continues to grow, it may become possible to train neural networks using experimental data instead of simulated data. Our research used simulated data to train our network, but experimental training data may improve results further. This change would prevent the model from over-fitting the simulated data and give it a wider, more representative data set to train with. Also, by adding an additional 1×1 scalar input to the network to denote the resolution of an experimental map, the C-CNN could train to differentiate between map resolution. This change would allow the network to learn each resolution with more independence.

This research relied on a manually chosen threshold value to normalize each experimental density map before evaluating it with the C-CNN. This manual step is not ideal as it requires subjective input by the user. Further improvements could remove this manual step by adding an automatic method which use traits such as map surface area or resolution to automatically calculate a threshold value for each density map. Another effective method could train a CNN, that also uses deep learning semantic segmentation, to automatically modifies density maps into a preprocessed state.

## V. CONCLUSION

In summary, we presented an effective method for protein backbone prediction from high resolution cryo-EM density maps using deep learning. This approach used three cascaded convolutional neural networks to produce confidence maps for some of the major structural components of proteins. These confidence maps were processed using a variety of novel method including a tabu-search path-walking algorithm to construct backbone traces and a helix-refinement step to improve the structure of α-helices. Additionally, a new protein mapping algorithm was used to build up full atomic models from two of the final prediction maps (EMDB-5778 and EMDB-8410). Our method out-performed the Phenix based fully automatic model building method by producing backbone traces that were more complete (88.5% vs. 66.8%) as measured by percentage of matching Cα atoms. Further research may improve this research field by incorporating other structural aspects of protein molecules within the cascaded convolutional neural network or training the networks with experimental data.

## Footnotes

1 https://www.rcsb.org/

2 Approximately 6000 unique maps were used for training. Each was rotated by 0°, 90°, 180° and 270° to increase the training data size by 4x.

3 A cutoff threshold was selected for each simulated resolution. At each resolution all voxels less than the threshold were set to zero in order to remove low intensity noise from the density map.

4 The manually selected threshold value was determined by viewing the density map in Chimera and selecting a cutoff value that made the SSEs appear similar in size and structure to the SSEs of the simulated density maps that were used to train the neural networks.

5 https://www.emdataresource.org/index.html

6 https://docs.scipy.org/doc/scipy/reference/generated/scipy.optimize.minimize.html

7 https://www.emdataresource.org/

